# Positional information and information flows in dynamic tissues

**DOI:** 10.64898/2026.04.15.718553

**Authors:** Alex Plum, Mattia Serra

## Abstract

During development, embryos store, transmit, and transform information to generate spatial patterns. Positional information (PI) quantifies how precisely cells form patterns at a given time, but cell motion has limited its application to static tissues. We introduce a framework for PI in dynamic tissues by decomposing mutual information between cells’ positions and properties over time into information flows contributing to PI preservation, loss and generation. These reveal information-theoretic signatures of ubiquitous developmental processes, including instruction, sorting and mixing, directly from data. Applying this framework to whole-embryo cell trajectories in *Drosophila*, mouse and zebrafish gastrulation, we provide local and global information-theoretic quantification of cell mixing and derive bounds on PI preservation imposed by tissue dynamics. Analyzing tissue flows as dynamical systems, we further show that morphogenesis structures mixing, preferentially preserving specific patterns. Finally, we derive inequality conditions for tracing generated PI to candidate information sources and distinguishing among alternative pattern-formation mechanisms, from programmed extracellular cues to self-organizing intercellular interactions.

## Introduction

Development transforms genetic information into an organism’s phenotype. At intermediate developmental stages, positional information (PI) quantifies the information available to cells about their positions and encoded in multicellular patterns. Introduced through Wolpert’s French Flag problem [1] and later formalized using information theory [2, 3], PI has been quantified in systems with approximately static cell positions, including the pre-gastrulation *Drosophila* embryo [2, 4, 5], the wing imaginal disc [6], and the vertebrate neural tube [7]. However, cell motion is ubiquitous during development, and PI has not yet been generalized to dynamic tissues undergoing cell rearrangement, proliferation, and morphogenesis [7–11], substantially limiting its applicability.

Bridging this gap is essential for understanding how embryos generate precise spatial patterns amid ongoing cell motion. Here, we introduce a general framework for PI in dynamic tissues. We treat embryos as stochastic dynamical systems and use information theory to elucidate how the dynamics of cell motion and cell properties affect PI over time. Because information theory provides a mechanism-independent quantification of statistical dependencies [12–15], this perspective offers a unifying language for diverse developmental processes and data modalities [3], linking tissue kinematics and underlying mechanisms to information-processing constraints.

In Section 1, we develop a finite-time framework for dynamic PI by decomposing mutual information between cells’ positions and properties into interpretable contributions to PI loss and generation, illustrated with minimal models. In Section 2, we investigate PI preservation using whole-embryo cell trajectory data across species. Treating dynamic tissues as information channels, we derive bounds on PI preservation imposed by cell mixing and uncover constraints between morphogenesis and patterning, biasing which patterns are easiest to preserve, with implications for fate maps. In Section 3, we characterize PI generation by tracing information flows to extracellular and intercellular information sources, enabling discrimination among candidate patterning mechanisms from data.

## 1 Decomposing positional information over time

Mutual information [12] formalizes PI as the information that cell positions (**x**, where cells are) provide about cell properties (**g**, their states) and vice versa [2]. Here, **x** and **g** are random variables with realizations ***x*** ∈ 𝒳 and ***g*** ∈ 𝒢 drawn from a joint probability distribution *p*(***x, g***). Positions ***x*** are sampled from physical space (i.e., 𝒳 is a set of 1D, 2D or 3D positions) while 𝒢 is a set of cell properties, with gene expression levels being a canonical choice. The positional distribution *p*(***x***) is often taken to be uniform over the discretized embryonic domain 𝒳 [2, 3, 6, 7], implying maximum entropy *H*(**x**) = log_2_ |𝒳| (|𝒳 | denotes the number of elements in 𝒳), although this assumption can be relaxed. The entropy *H*(**g**) = − ∑_***g***∈G_ *p*(***g***) log_2_ *p*(***g***) (log = log_2_ hereafter) quantifies variability in cell properties. Computed from *p*(***x, g***), PI is the mutual information (MI) between positions and properties:

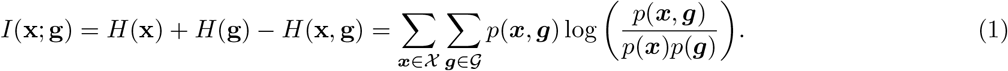

It is helpful to visualize information-theoretic quantities with I-diagrams, analogous to Venn diagrams (Fig. 1). In I-diagrams, entropies *H*(**x**) and *H*(**g**) correspond to the areas of regions, here disks 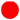 and 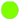. Unions 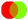 correspond to joint entropies *H*(**x, g**), differences 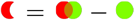 to conditioning *H*(**x**|**g**) = *H*(**x, g**) − *H*(**g**), and intersections 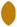 measure *I*(**x**; **g**) [16]. To study information dynamics over a time interval [0, *t*]—e.g. a developmental stage—we consider **x**_0_, **g**_0_, **x**_*t*_, and **g**_*t*_ (Fig. 1A), distinguishing initial PI *I*(**x**_0_; **g**_0_) 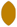 from final PI *I*(**x**_*t*_; **g**_*t*_) 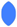 (Fig. 1B). This finite-time view introduces new MI terms between **x, g** over time (Fig. 1C). For example, *I*(**x**_0_; **x**_*t*_) 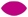 and *I*(**g**_0_; **g**_*t*_) 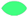 are both examples of finite-time *predictive information* [17, 18], quantifying how well cells’ initial positions and properties predict their later positions and properties. During development, both *H*(**x**) and *H*(**g**) typically increase as proliferating and differentiating cells expand the ranges of positions and properties (|𝒳|, |𝒢|) (Fig. 1C). To generate increasingly complex patterns during development, properties and positions must increase their MI. In addition to *I*(**x**_0_; **x**_*t*_) and *I*(**g**_0_; **g**_*t*_), our framework introduces additional *information atoms* 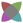 [16] marked by regions where three or more variables overlap (Fig. 1A) and corresponding to PI preserved, lost, or generated in different ways (Fig. 1D).

**Fig. 1:**
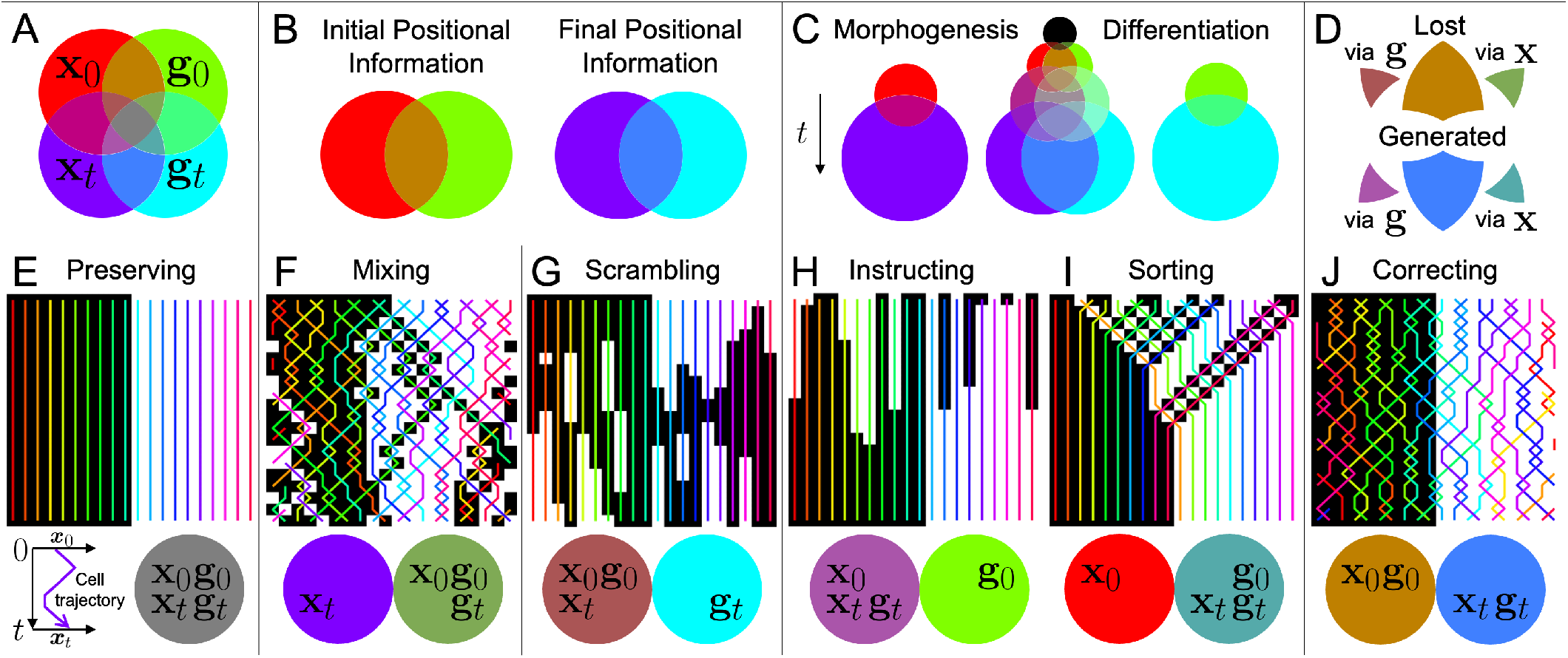
Decomposition of positional information over time. A) Four-variable I-diagrams for initial positions **x**_0_ 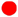, initial properties **g**_0_ 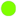, final positions **x**_*t*_ 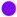, and final properties **g**_*t*_ 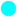. Overlapping disks average their colors. Each color identifies a distinct *information atom*. The area of each disk represents the Shannon entropy (i.e., uncertainty) of the corresponding random variable. B) Positional information (PI) is the overlap area between positions and properties at a given time. C) Development tends to increase positional entropy through morphogenesis 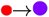 and property entropy through differentiation 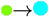. An expanding I-diagram illustrates how embryos develop more complex patterns by increasing PI over successive developmental stages. Overlaps between initial and final variables 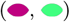 quantify predictive information for positions and properties, i.e., how well initial states (**x**_0_ or **g**_0_) predict final states (**x**_*t*_ or **g**_*t*_). D) Changes in PI decompose into PI lost (terms 2-4 of Eq. (2)) and PI generated (terms 5-7). E-J) Minimal discrete 1D models isolating terms in Eq. (2). Top: Representative simulations show cell trajectories (colored lines) and binary cell properties (black/white) over time (top to bottom). Bottom: I-diagrams, where a full disk represents one bit. E) Perfect preservation. F) Cell position mixing loses PI. G) Scrambling (i.e., unpredictability in cell property dynamics) loses PI. H) Instructing generates PI via information flow from **x**_0_ → **g**_*t*_. I) Sorting generates PI via information flow from **g**_0_ → **x**_*t*_. J) Correcting compensates for PI losses via information flows from **x** to **g** along trajectories. See Sections S1.2-S1.3 for model details and Fig. S1 for time series of the terms in Eq. (2) for each model.

To understand PI dynamics, we algebraically decompose final PI into initial PI and terms corresponding to PI lost and generated (Section S1.1). PI loss and generation can be captured by conditional mutual information (CMI) terms, which measure the information shared by two variables that remains after conditioning on other variables:

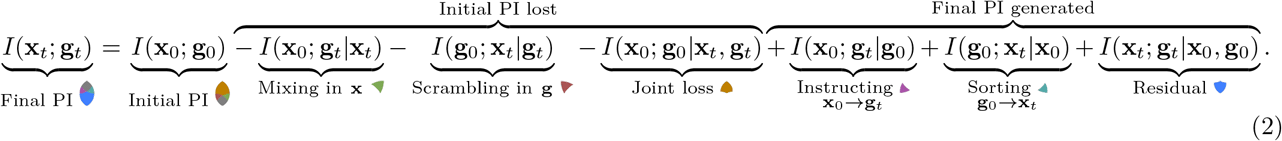

Terms 1 − 4 of Eq. (2) quantify preserved PI 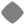—the overlap between initial and final PI—by subtracting PI losses from initial PI 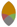. PI losses include atoms where all variables overlap except **x**_*t*_ 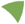 or **g**_*t*_ 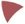 and where only **x**_0_ and **g**_0_ overlap 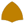 (Fig. 1A). Terms 5 − 7, instead, quantify generated PI to obtain final PI 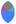. PI generation includes atoms where all variables overlap except **x**_0_ 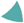 or **g**_0_ 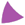 and where only **x**_*t*_ and **g**_*t*_ overlap 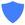 (Fig. 1D). Eq. (2) holds in general, derived from the chain rule for MI [19]. To gain intuition for the terms in Eq. (2), we consider six minimal models of prototypical developmental processes (Fig. 1E–J). For illustration, we restrict **x** and **g** to two equiprobable realizations: cells are left or right (𝒳 = {L, R}) and black or white (𝒢 = {B, W}). This limits PI to 0 − 1 bit and yields I-diagrams with equal-sized disks. Interpretations of terms in Eq. (2) as lost or generated PI can be confounded by synergy [16, 20] (Section S1.4), which is absent in the examples below.

Models in Fig. 1E–G illustrate PI preservation and loss, and involve only the first three terms in Eq. (2). If neither **x** nor **g** changes, all of 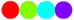 overlap as 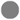, preserving *I*(**x**_0_; **g**_0_) = 1 bit (Fig. 1E). In dynamic tissues, positions evolve with nontrivial dependencies characterized by the probability *p*(***x***_*t*_|***x***_0_) that a cell ends at ***x***_*t*_ given that it started at ***x***_0_ (Fig. 2E), possibly leading to PI loss. If motion is dominated by cell mixing, it may erase the initial pattern (Fig. 1F). In this case, properties are fixed, and the PI lost over time is *I*(**x**_0_; **g**_*t*_|**x**_*t*_) 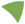, which increases to 1 as *I*(**x**_0_; **x**_*t*_) decreases to 0. The term *I*(**x**_0_; **g**_*t*_|**x**_*t*_), therefore, quantifies how cell mixing deteriorates initial PI and is related to the predictive information between initial and final cell positions *I*(**x**_0_; **x**_*t*_) (Section 2). Similarly, in Fig. 1G, stochastic cell-property changes spread out *p*(***g***_*t*_|***g***_0_). This unpredictability in property dynamics, which we term *scrambling* to distinguish it from *positional mixing*, likewise erases the initial pattern. In this case, positions are fixed and the loss is quantified by *I*(**g**_0_; **x**_*t*_|**g**_*t*_) 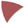, which rises to 1 as *I*(**g**_0_; **g**_*t*_) falls to 0. In Section 2, we use these loss terms to show how mixing constrains pattern preservation.

**Fig. 2:**
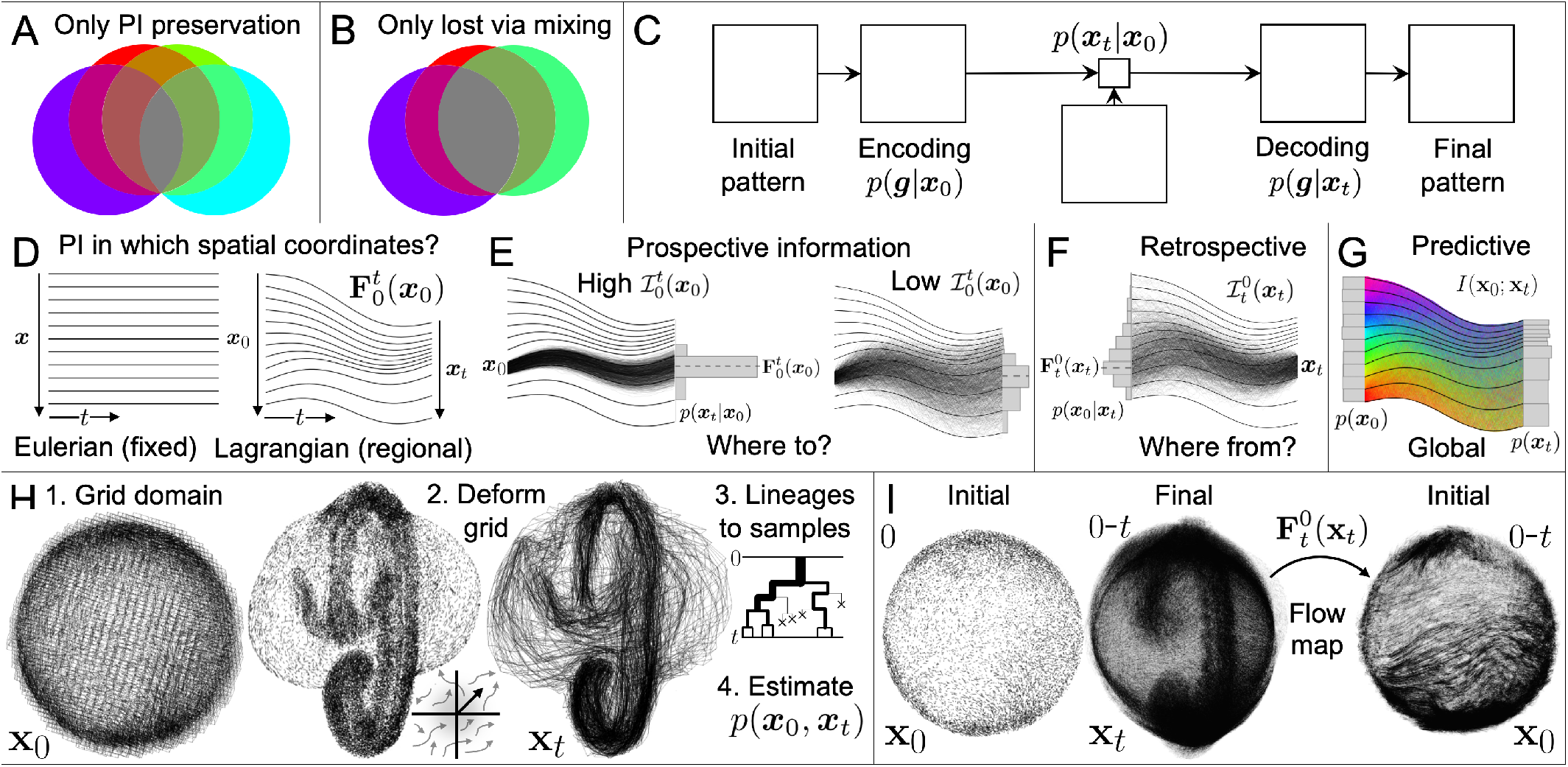
Preserving positional information. A) Four-variable I-diagram with no PI generation described by Eq. (3). Colors as in Fig. 1. B) I-diagram further fixing properties **g**_*t*_ = **g**_0_, so that PI is lost only through cell mixing. C) Information channel for preserving PI in dynamic tissues (Eq. (4)). An initial pattern, encoded by *p*(**g**|**x**_0_), is transmitted through noisy positional dynamics *p*(***x***_*t*_|***x***_0_), and decoded from the final positions *p*(**g**|***x***_*t*_). D) Eulerian coordinates track fixed spatial locations vs Lagrangian coordinates labeling tissue regions that move with the tissue, represented by the flow map 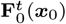. E) Prospective information 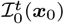 (Eq. (5a)) quantifies how well specific initial cell positions predict final positions. High (low) prospective information indicates low (high) mixing. F) Retrospective information 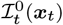 quantifies how well specific final cell positions predict initial positions. G) Predictive information *I*(**x**_0_; **x**_*t*_) is obtained by averaging prospective or retrospective information over all regions, providing a global measure of cell mixing (Eq. (5b)). H) Pipeline for estimating information-theoretic quantities from whole-embryo cell trajectory data. I) Because the flow map—describing average tissue motion—is invertible, trajectories can be visualized either on the final configuration or mapped back onto the initial, undeformed configuration.

Models in Fig. 1H–J illustrate PI generation through the final three terms in Eq. (2). Crucially, all three require dependent dynamics between **x** and **g**. In Fig. 1H, cells’ properties are instructed according to their fixed positions, as in Wolpert’s gradient model [1]. The CMI *I*(**x**_0_; **g**_*t*_|**g**_0_) 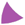 quantifies an *information flow* [21, 22] from **x**_0_ to **g**_*t*_, namely the additional information that initial positions provide about final properties beyond initial properties alone. While static tissues permit only position-to-property information flows, dynamic tissues also permit property-to-position information flows. In Fig. 1I, cells sort according to their fixed properties, and *I*(**g**_0_; **x**_*t*_|**x**_0_) 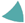 quantifies the information flow from **g**_0_ to **x**_*t*_. Finally, in Fig. 1J, PI is preserved despite losses of predictive information in both **x** and **g**, captured by *I*(**x**_0_; **g**_0_|**x**_*t*_, **g**_*t*_) 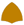. These joint losses are offset by continual error correction—coordinated changes in **x**_*t*_ and **g**_*t*_ along trajectories (Section S3.2)—generating PI not traceable to **x**_0_ or **g**_0_, and therefore captured by *I*(**x**_*t*_; **g**_*t*_|**x**_0_, **g**_0_) 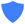. In Section 3, we trace generated PI to latent information sources—extracellular cues and intercellular interactions—that mediate these information flows.

Mutual information is an integrated (scalar) quantity that averages over all possible positions and properties (Eq. (1)). It is informative, however, to elucidate *which* properties and positions i) share information, ii) lose predictability, and iii) interact dynamically to generate precise patterns. This local information structure is captured by *pointwise mutual information* (PMI) [23]. The PMI between pairs of realizations is *i*(***x***; ***g***) = log (*p*(***x, g***)*/*(*p*(***x***) *p*(***g***))), measuring how much more (or less) likely ***x*** and ***g*** are to co-occur than if they were statistically independent. Mutual information (Eq. (1)) is the expected PMI, *I*(**x**; **g**) = ∑_***x***_ ∑_***g***_*p*(***x, g***) *i*(***x***; ***g***). To gain intuition about PMI, consider the examples in Fig. 1E-J, where *p*(***x***) = *p*(***g***) = 1*/*2. With no pattern, all pairs are independent and *i*(***x***; ***g***) = 0 because *p*(***x, g***) = *p*(***x***)*p*(***g***) = 1*/*4. With a perfect pattern, only *i*(L, B) and *i*(R, W) contribute to *I*(**x**; **g**) because *p*(L, B) = *p*(R, W) = 1*/*2 while *p*(L, W) = *p*(R, B) = 0. For Fig. 1H, pointwise CMI *i*(***x***_0_; ***g***_*t*_|***g***_0_) confirms that ***x***_0_ = L (R) predicts ***g***_*t*_ = B (W), while for Fig. 1I, *i*(***g***_0_; ***x***_*t*|_***x***_0_) confirms that ***g***_0_ = B (W) predicts ***x***_*t*_ = L (R). In Sections 2.2-2.5, we leverage PMI to localize information dynamics in space. More generally, pointwise forms of MI and CMI help to connect global information dynamics—PI over time—to specific spatial locations and underlying mechanisms. We summarize the information-theoretic quantities used throughout in Table 1.

**Table 1:**
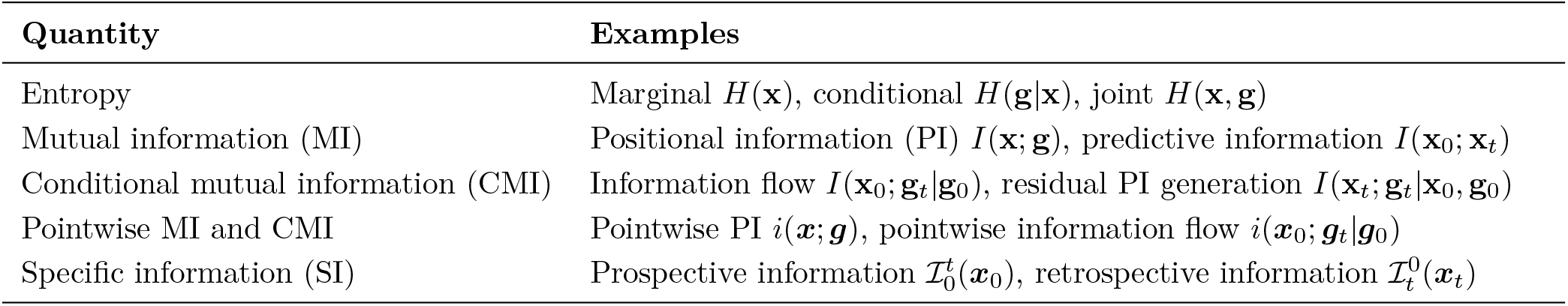
Distinct information-theoretic quantities with representative examples.

## 2 Preserving positional information

Mixing and scrambling decrease PI and constrain pattern preservation. To isolate these constraints, we first neglect PI generation. If position and property dynamics are independent, with **x**_*t*_ depending only on **x**_0_ and **g**_*t*_ depending only on **g**_0_, then **x**_*t*_ and **g**_*t*_ are conditionally independent given either **x**_0_ or **g**_0_. This eliminates all PI generation terms in Eq. (2), leaving only PI preservation, which can be rewritten (Section S2.1) as:

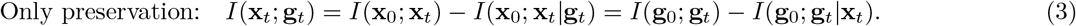

Fig. 2A shows a representative I-diagram for Eq. (3), distinct from Fig. 1A in that it lacks the information atoms for PI generation (Fig. 1D). The predictive information *I*(**x**_0_; **x**_*t*_)—our focus in this section—quantifies the fidelity with which initial positions predict final positions, maximized when **x**_0_ determines **x**_*t*_ (Fig. 1E,G,H) and minimized when they become independent (Fig. 1F,I,J). The subtracted terms in Eq. (3) account for mixing in **x** and scrambling in **g** that do not alter final PI—e.g., the mixing of cells with the same properties. Without PI generation, Eq. (3) shows that final PI is upper-bounded not only by *I*(**x**_0_; **g**_0_) (Eq. (2)), but also by the predictive information in **x** and **g**, independent of the initial pattern.

### 2.1 Dynamic tissues as channels for PI

Systems limited to preserving—but not generating—information can be formalized as information channels. Shannon defined the channel capacity *C* as the maximum mutual information achievable through a channel over all possible input encodings [12]. If only **x** changes (Fig. 2B)—a good approximation when positional dynamics are faster than cell differentiation—then the positional dynamics *p*(***x***_*t*_|***x***_0_) define a channel through which PI is transmitted (Fig. 2C), with prepatterns encoded by *p*(***g***|***x***_0_) and losses due solely to positional mixing (term 2 of Eq. (2), Section S2.2). Using Eq. (3), we derive the channel capacity of the dynamic tissue:

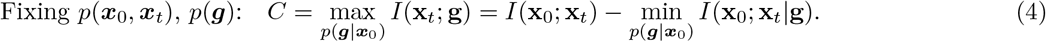

This capacity depends on i) *I*(**x**_0_; **x**_*t*_) and ii) the minimum *I*(**x**_0_; **x**_*t*_|**g**) achievable by prepatterns—for example, by assigning identical properties to adjacent cells prone to exchanging positions (Fig. 1F). Previous work has identified channel capacities at the cellular scale, constraining information transmission from external signals to gene expression [24], in gene-regulatory elements [25], and from experimental perturbations to cell type proportions [26]. Eq. (4) instead constrains the embryo-scale transmission of spatial patterns. Notably, this upper bound, *I*(**x**_0_; **x**_*t*_), depends only on cell trajectories, which are increasingly accessible in modern datasets [27]. Moreover, position is the common currency of all PI, so its dynamics constrain patterning precision in any cell property. *I*(**x**_0_; **x**_*t*_) provides the nonlinear generalization [28] of linear correlation previously used to analyze positional dynamics in specific regions [9,29]. While *I*(**x**_0_; **x**_*t*_) has been used to quantify mixing in fluids [30], it remains unexplored in developmental biology. To estimate *I*(**x**_0_; **x**_*t*_) from cell tracks, we first discuss the appropriate coordinates to represent positions in dynamic tissues.

### 2.2 Eulerian vs. Lagrangian spatial coordinates

Fig. 1F illustrates how mixing loses PI in a minimal 1D model with a fixed domain. In embryos, however, cells move within dynamic, deforming tissues in 3D. A core challenge in extending PI to dynamic tissues is how to quantify and compare positions over time [15]. In static tissues, ***x*** may represent an absolute distance from a landmark or a position rescaled relative to multiple landmarks [5]. In dynamic tissues, however, these landmarks move [7], and different coordinate choices—e.g., absolute versus rescaled positions—yield different PI estimates [5]. Thus, the definition of ***x***_0_ and ***x***_*t*_ should 1) require minimal alignment as embryos drift or rotate, and 2) adapt consistently with tissue deformations. The fixed Eulerian coordinates ***x*** = [*x, y, z*] in which trajectory data are typically recorded lack both features. Instead, we consider Lagrangian coordinates (Fig. 2D), which label material regions that move and deform with the embryo, including boundaries and internal landmarks. The *flow map* 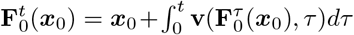 describes the average motion of regions in a given embryo, where ***x***_0_ labels regions’ initial positions and 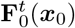 their final positions (Fig. 2D). With cell mixing (in addition to average motion), cells starting in ***x***_0_ may end in different regions 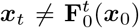, and the conditional probabilities *p*(***x***_*t*_|***x***_0_) quantify these transitions (Fig. 2E). Transforming Eulerian to Lagrangian coordinates reframes position from “Where in space?” to “Which region?”, consistent with the regional labels used in fate mapping [31].

### 2.3 Local and global measures of cell mixing

Reliable fate maps require consistent positional mappings (low mixing) between initial and final regions [31–33], and this fidelity is often spatially heterogeneous. Fate maps can be *prospective*—following cells forward in time—and visualizing their eventual fates on the embryo’s initial configuration, or *retrospective*—following cells back to where they came from [31]. By quantifying *specific information* (SI) that a position provides about its past or future, we can similarly assess local information dynamics and pinpoint where positional mappings retain precision. For example, considering the final regions ***x***_*t*_ that cells originating in a given initial region ***x***_0_ may reach, the prospective information 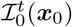 quantifies how predictive a specific ***x***_0_ is about ***x***_*t*_ (Eq. (5a)):

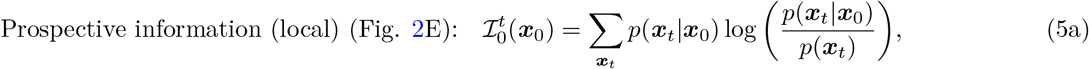

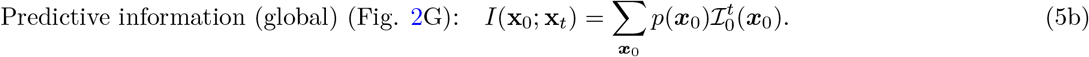

Prospective information measures the divergence between *p*(***x***_*t*_|***x***_0_) and *p*(***x***_*t*_)—a nonnegative number of bits. The expectation of 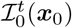 recovers the predictive information (Eq. (5b))—a global measure of cell mixing. Initially, 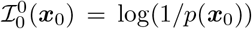 is maximal as cells have not moved. This information is lost as *p*(***x***_*t*_|***x***_0_) spreads over 𝒳, reaching 0 if it becomes identical to *p*(***x***_*t*_) (e.g., uniform). While *i*(***x***_0_; ***x***_*t*_) = log (*p*(***x***_0_, ***x***_*t*_)*/*(*p*(***x***_0_)*p*(***x***_*t*_))) (PMI) and *I*(**x**_0_; **x**_*t*_) (MI) are symmetric, 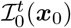 is forward-looking. Its backward-looking dual, *retrospective information* 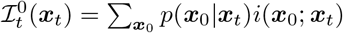 quantifies information about where cells in a specific ***x***_*t*_ came from (Fig. 2F).

### 2.4 Quantification of whole-embryo cell mixing across species

We apply our framework to whole-embryo cell trajectory data [34] from *Drosophila* [35], zebrafish [36,37] and mouse [38] gastrulation. Estimating probabilities typically requires many embryos (a developmental ensemble [3]), but in estimating *p*(***x***_0_, ***x***_*t*_), multiple cells occupying the same discrete region enable probability estimates from a single embryo by trading spatial resolution for sample size. For example, each model realization in Fig. 1E-J yields 10 samples per L*/*R region. These datasets contain roughly 5, 000–25, 000 cells per time point with temporal resolutions of 30 seconds to a few minutes [35–38]. Previous studies have assessed mixing in specific tissues using trajectory statistics, such as mean squared displacement, by subtracting mean tissue motion or fitting it to distinguish random from directed motion [39, 40]. These approaches rely on ad hoc choices and cannot accommodate the space-dependent mean motion ubiquitous in whole embryos [41]. Moreover, averaging relative displacements between initially neighboring cells [42] does not distinguish stochastic rearrangements from deterministic deformation. Our approach resolves these limitations by incorporating mean motion into Lagrangian coordinates and explicitly capturing inter-region mixing.

To estimate information-theoretic quantities from trajectory data, we: i) grid the initial domain at a chosen spatial resolution, balancing spatial resolution against the number of samples (i.e. cells) per region; ii) infer the deformation of this three-dimensional grid over time from cell trajectories to obtain a coarse 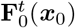 describing the mean tissue motion; iii) convert lineage data (mother-daughter relationships) into distinct samples {***x***_0_, ***x***_*t*_*}* from trajectories present at *t* and traceable back to time 0, so that dividing cells can contribute multiple samples while excluding cells that appear after 0 or disappear before *t*; and iv) count samples initially occupying ***x***_0_ and finally occupying ***x***_*t*_ to construct 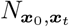. We then estimate joint 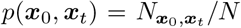, marginal (*p*(***x***_0_),*p*(***x***_*t*_)) and conditional (*p*(***x***_0_|***x***_*t*_),*p*(***x***_0_|***x***_*t*_)) probabilities (Fig. 2H), where 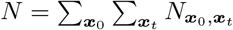 is the total number of samples. The flow map can be inverted to visualize trajectories either in the final configuration or mapped back to the initial configuration (Fig. 2I). Sections S1.3 and S2.3 provide methodological details. Fig. 3A-C provide the first information-theoretic quantification of mixing in embryos, revealing mixing patterns directly from trajectory data. Each embryo begins gastrulation (Fig. 3A-C, left, **x**_0_) with approximately uniform prospective information (Eq. (5a)). This information is lost as cells mix, visualized on the final configurations (***x***_*t*_, Fig. 3A-C, middle and right) or on the initial configuration (***x***_0_, Fig. 3F, right).

**Fig. 3:**
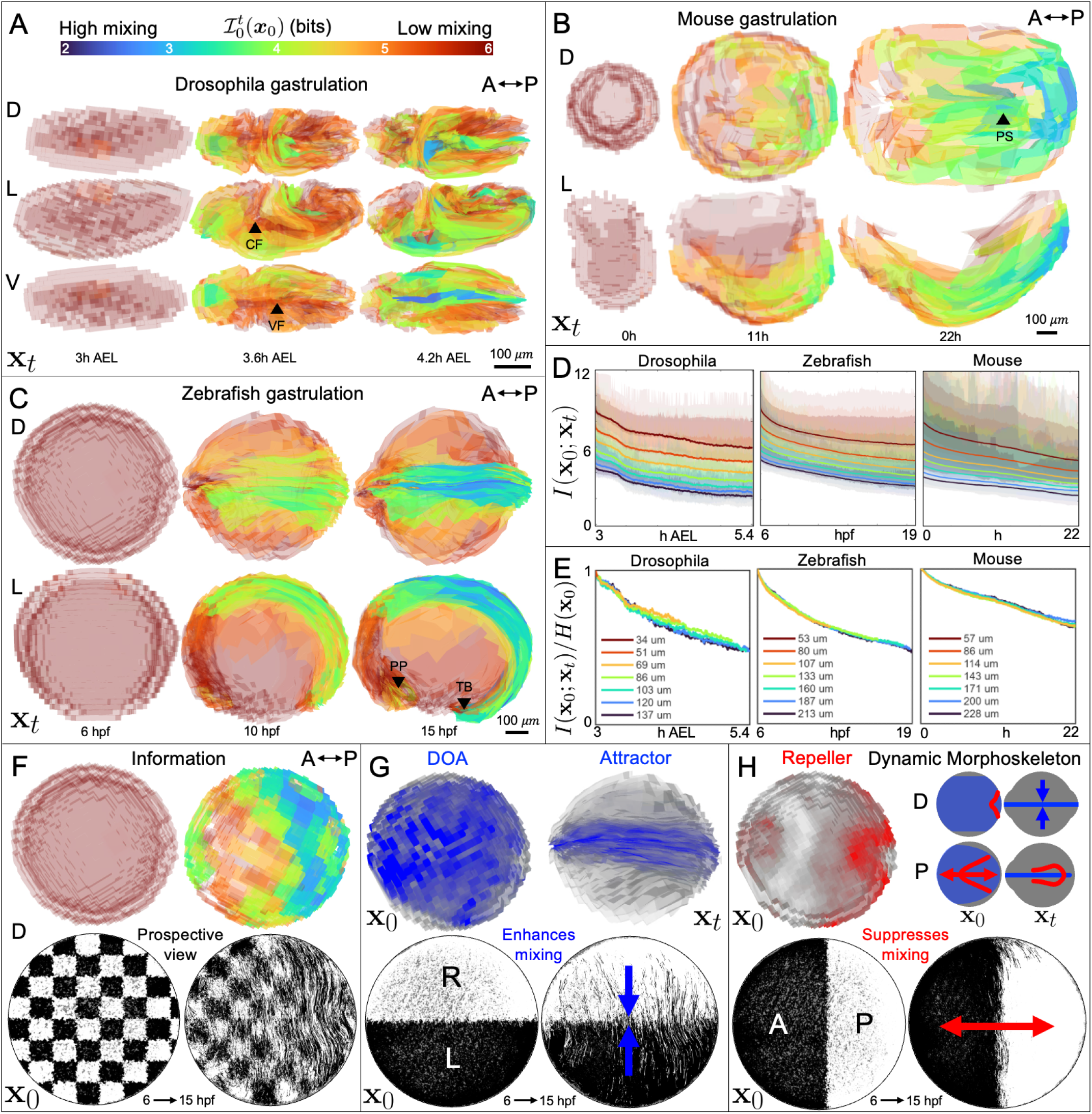
Whole-embryo quantification of cell mixing during gastrulation. A-C) Prospective information 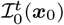 (Eq. (5a)), estimated over time from whole-embryo trajectories in *Drosophila*, mouse and zebrafish gastrulation. We visualize 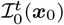 on embryos’ deformed configurations ***x***_*t*_ to associate 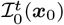 with morphological structures of interest, including the cephalic furrow (CF), ventral furrow (VF), primitive streak (PS), prechordal plate (PP) and tailbud (TB). View labels: D, dorsal; L, lateral; V, ventral. Low-mixing regions (red) are rendered more transparent for 3D visualization. D) Predictive information *I*(**x**_0_; **x**_*t*_) (Eq. (5b)) over time at different spatial resolutions (solid lines). Shaded bands show the 5 − 95% range of prospective information across regions. E) Finite-sample-corrected estimates of *I*(**x**_0_; **x**_*t*_)—shown only for resolutions and times for which shuffling the data leaves negligible residual (bias-induced) information (Section S2.4)—normalized by their initial values *H*(**x**_0_). F) 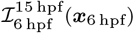 in zebrafish indicates spatial heterogeneity in mixing. Bottom: Zebrafish cell trajectories mapped onto the initial configuration via the inverse mean motion illustrate heterogeneous degradation of a checkered prepattern, as well as directional mixing (deterministic cell motion would leave the checkered pattern unchanged). G-H) Zebrafish dynamic morphoskeleton [43] for 6-10hpf, consisting of attractors (blue) and repellers (red) shown on the initial (**x**_0_) and final (**x**_*t*_) embryo configurations (Section S2.5). Bottom: Sharp binary prepatterns degrading as zebrafish cell trajectories mix, visualized on the initial configuration. G) Mediolateral (L-R) pattern degrades more as the domain of attraction (DOA) (left) converges to the attractor (right). H) Anteroposterior pattern preferentially preserved by extension along the A-P axis, associated with a repeller. Schematics depict dorsal and posterior views of the DOA, attractor, and repeller (dynamic morphoskeleton) on the initial and final configuration.

In the *Drosophila* embryo, mixing accompanies germband extension via cell intercalation [44,45], whereas the cephalic and ventral furrows form without extensive rearrangement. Consistent with this, Fig. 3A shows initially low mixing in both furrows, followed by high mixing of spreading mesoderm after ventral furrow collapse [46]. In the mouse embryo, cells rearrange during convergence toward the primitive streak and individually ingress [47]. Retrospective fate mapping—tracing labeled cells back to ***x***_0_— [38] and motility quantification [48] suggest pronounced mixing in the posterior, consistent with the low posterior prospective information in Fig. 3B. In the zebrafish embryo, Fig. 3C shows substantial mixing in the posterior body axis, where mesoderm predominates, whereas the anterior axis and lateral regions retain prospective information. Anterior mixing is pronounced only beneath the prospective forebrain, where prechordal-plate mesoderm cells delaminate and are more motile than the surrounding ectoderm [49]. These results show that information theory reveals biologically meaningful mixing patterns directly from cell trajectory data.

To quantify how mixing patterns constrain PI preservation, we estimate *I*(**x**_0_; **x**_*t*_) (Eq. (5b)), which upper-bounds the PI channel capacity (Eq. (4)). Fig. 3D shows *I*(**x**_0_; **x**_*t*_) over time across spatial resolutions. At a given resolution, *I*(**x**_0_; **x**_*t*_) constrains PI preservation for patterns at that resolution (Eq. (3)). Finer resolutions make more positional distinctions and therefore begin with more predictive information to lose, but they also provide fewer samples for estimation. Because we require trajectories present at both the initial and final times, sample sizes also decrease over time. To assess the reliability of *I*(**x**_0_; **x**_*t*_) estimates and their dependence on spatial resolution, we correct the plug-in estimates in Fig. 3D for finite-sample bias, exclude estimates that fail a shuffle test (Section S2.4), and normalize the remaining curves by their initial values for comparison (Fig. 3E). In all three species, the normalized curves approximately collapse across resolutions. Remarkably, all three species lose approximately 40% − 50% of their predictive information during gastrulation, despite differences in size and duration. With two regions, for example, this corresponds to roughly 10% of cells exchanging regions.

Prospective information captures heterogeneity in mixing, but not directionality, and it cannot determine *which patterns* minimize the loss in Eq. (4). Fate patterns often comprise discrete compartments delimited by stably inherited boundaries [50]. Only mixing across these boundaries degrades PI. For example, in Fig. 1F, PI decreases as cells mix across the L*/*R boundary. In 2D and 3D settings, mixing directions may be regulated to support compartment boundaries. We investigate this effect using zebrafish cell trajectory data, which provide the highest cell counts and simplest, initially spherical geometry. To isolate mixing from morphogenesis, we map all trajectories back to this initial configuration via the inverse flow map (Fig. 2I), generalizing prior approaches that remove bulk motion by subtracting mean displacements [39]. Prospective information on the initial configuration (Fig. 3F, top) shows high mixing (low 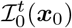) in the posterior, also apparent in how a checkered prepattern degrades (visualized on the initial configuration in Fig. 3F, bottom). Strikingly, trajectories on the initial configuration reveal pronounced medio-lateral mixing. Directional mixing may arise from structural barriers maintained by junctional tension [51, 52], or from intrinsically anisotropic random cell motility. Below, we show that the observed directional mixing (Fig. 3F) can be induced by morphogenesis itself, consistent with findings that morphogenesis mediates diffusive morphogen transport [53].

### 2.5 Morphogenesis mediates mixing

Cell rearrangement often accompanies morphogenesis, resulting in both directed and random neighbor exchanges. Pre-existing patterns must withstand the associated mixing to transmit a precise pattern [10]. Typically, cell motion can be decomposed into a deterministic velocity field (**v**(***x***, *t*), the regional-average motion driving morphogenesis) and a stochastic component (effective diffusivity **D**(***x***, *t*), mixing) [39, 42, 54]. This diffusivity may be nonuniform, non-constant, and anisotropic. To isolate directionality induced by morphogenesis, we take mixing to be isotropic **D**(***x***, *t*) = *D*(***x***, *t*)**I**. The conditional probability over Eulerian coordinates ***x*** (Fig. 2D), describing positions at time *t* for cells starting at (***x***_0_, *t* = 0) obeys, in the incompressible case (**∇ · v** = 0), the Fokker–Planck equation [55]:

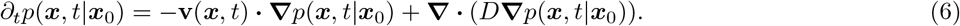

As we have seen, the Eulerian coordinates ***x*** in Eq. (6) are not the appropriate ones for PI dynamics, because the relevant question concerns mixing between deforming tissue regions. We therefore transform Eq. (6) into Lagrangian coordinates that move with the average tissue flow 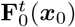, Fig. 2D). In these coordinates, with ***x***_0_, 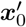 labeling regions in the initial configuration, the conditional probability evolves as

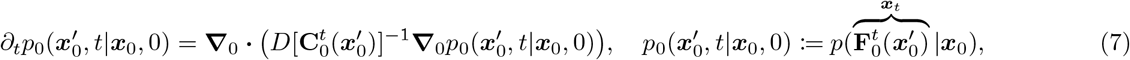

where *p*_0_ defines a probability density over the initial configuration [55, 56]. The Cauchy-Green strain tensor 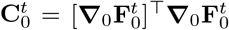 quantifies finite-time regional deformation using spatial gradients **∇**_0_ over the initial configuration [43]. See Section S2.6 for the expanded form of Eq. (7) with compressible flows and anisotropic **D**(***x***_0_, *t*). In morphogenesis without mixing (*D* = 0), cells move deterministically and 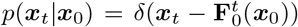. In mixing without morphogenesis, regions do not move relative to one another 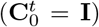 and mixing reduces to simple diffusion. With both morphogenesis and mixing present, Eq. (7) shows that morphogenesis mediates regional exchanges through 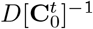. Eq. (7) therefore links information dynamics to the interplay between random cell motility (*D*) and morphogenesis 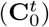 (Section S2.7). Notably, 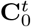 is frame-invariant, demonstrating that rigid rotation, translation, and other frame-dependent motion do not affect information dynamics.

We next examine how morphogenesis affects which patterns are most easily preserved, i.e., minimize *I*(**x**_0_; **x**_*t*_|**g**) (Eq. (4)). With uniform, isotropic *D* in a static tissue, the most transmissible patterns minimize compartment-boundary areas and therefore inter-compartment fluxes. On a symmetric domain such as a disc or sphere, these optimal patterns may be oriented arbitrarily. In a dynamic tissue, directional mixing—induced by anisotropic 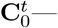 breaks this symmetry and biases pattern transmissibility. As a minimal model, consider linear convergent extension at rate *α*: tissue converges along a left-right (L-R) axis and extends along an anterior-posterior (A-P) axis. In this case, 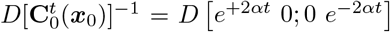 becomes directional, distinguishing L-R (+*αt*) and A-P, (−*αt*) directions, and causing L-R patterns to degrade faster than A-P patterns. Real morphogenetic movements are spatiotemporally heterogeneous. Lagrangian Coherent Structures, and specifically the *dynamic morphoskeleton* (DM) [43], identify where tissue regions maximally converge (*attractors*) or separate (*repellers*) from the eigenvalues of 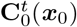 (Section S2.5). Eq. (7) implies that attractors are the strongest enhancers of mixing while repellers are the strongest barriers. In zebrafish, the DM includes an attractor along the forming body axis (Fig. 3G), intersected by a posterior repeller (Fig. 3H). As predicted by Eq. (7), Fig. 3G-H show that, medio-lateral prepatterns (e.g., L-R) degrade faster due to convergence to the attractor, whereas A-P prepatterns are preferentially preserved by axis elongation (repeller).

In *Drosophila*, directional mixing associated with convergent extension helps explain how dorsoventral intercalations preserve pair-rule gene stripes perpendicular to the extending germband [44]. In mouse, it may explain why track-pair crossings are less frequent along the A-P axis than the proximal-distal axis [48]. Similar associations between repellers and compartmentalized cell populations are observed in zebrafish [36] and chick [57]. Notably, many vertebrate embryos partition fate domains along the extending A–P axis, often prior to L–R symmetry breaking [58]. Together, these observations suggest an internal constraint [59, 60] between morphogenesis and pattern transmissibility, favoring prepattern orientations aligned with subsequent tissue flows. The agreement between information-theoretic measures of mixing, dynamical structures in the DM, and known fate maps suggests that embryos contend with this constraint.

## 3 Generating positional information

Overcoming constraints on PI preservation requires information flow from **x**_0_ to **g**_*t*_ (Fig. 1H, 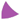), from **g**_0_ to **x**_*t*_ (Fig. 1I, 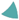), or along trajectories (Fig. 1J, 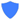), recovering terms 5-7 in Eq. (2). PI generation is more challenging to characterize than PI preservation because generated PI must be traceable to latent information sources in the embryo or its environment. Tracing PI amounts to determining i) how information flows between **x** and **g** (Eq. (2)), ii) which information sources are necessary from perturbation experiments, and iii) whether those sources supply sufficient information. This logic underlies PI studies in the pre-gastrulation *Drosophila* embryo, where minimal movement reduces the problem to a **x**_0_ → **g**_*t*_ flow. Maternal mutation experiments established Bicoid as a necessary information source [61]. Subsequent studies showed that although the Bicoid concentration profile is precise [62] and carries substantial positional information [2], its local concentration alone is insufficient to account for PI in the downstream gap genes [2,4], implying additional information sources. Here, we generalize this logic to dynamic tissues, discriminating among pattern-forming mechanisms in which cell movement may also be implicated.

### 3.1 Tracing PI generation to information sources

Whether information flows from **x** → **g**, from **g** → **x**, or bidirectionally to generate PI is not always clear. In 3D gastruloids, for example, both instructing and sorting mechanisms could break A-P symmetry. Early unpatterned Wnt activity (***g***_0_) predicts final axial position (***x***_*t*_), suggesting sorting mechanisms [63]. However, because both **x** and **g** change, the possibility of accompanying instructive mechanisms is difficult to exclude [63, 64]. When both **x** and **g** change [63–65], the initial states **x**_0_, **g**_0_ need not be informative, leaving residual PI generation *I*(**x**_*t*_; **g**_*t*_|**x**_0_, **g**_0_) in Eq. (2) and implying flows between **x** and **g** along trajectories (Fig. 1J). This residual PI generation can be eliminated by using trajectory-aware decompositions of *I*(**x**_*t*_; **g**_*t*_) involving 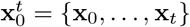 or 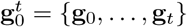 (Section S3.2).

Tracing PI simplifies if only one variable is dynamic (Fig. 1H-I). If either **x** or **g** is fixed, PI changes depend only on the dynamic variable, and residual PI generation vanishes. In these cases, Eq. (2) simplifies to (Fig. 4A-B):

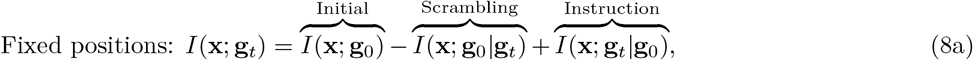

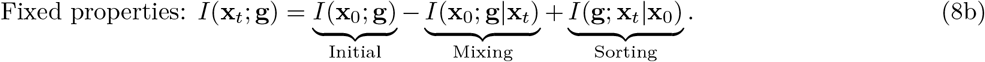

**Fig. 4:**
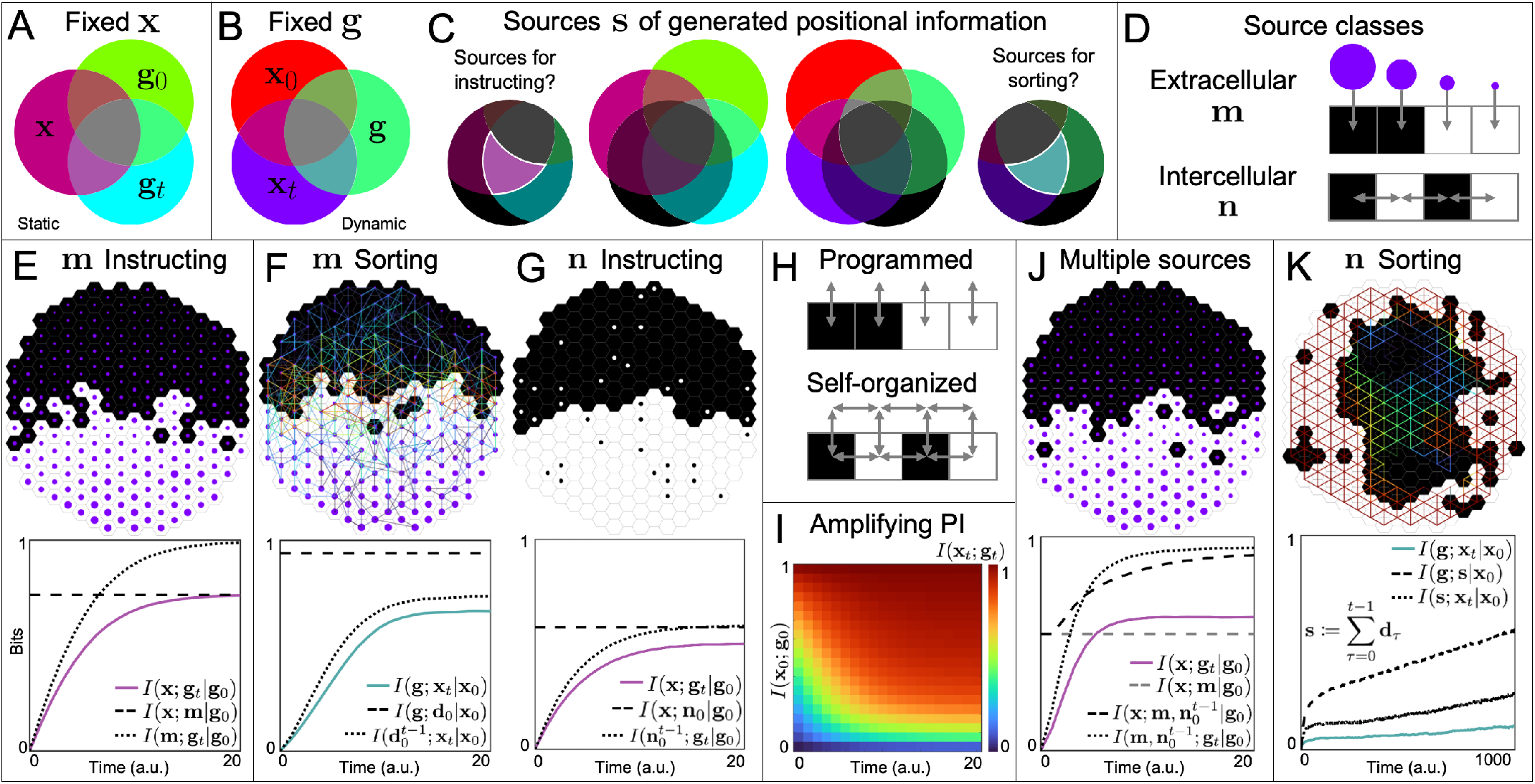
Generating positional information. A) I-diagram with fixed positions: PI can increase through property dynamics (instruction 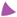, Eq. (8a)). B) I-diagram with fixed properties: PI can increase through positional dynamics (sorting 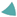, Eq. (8b)). C) I-diagrams for testing whether a candidate information source **s** (black disk) is sufficient to account for PI-generating information flows (Eq. (9)). D) Broad source classes: extracellular cues (**s** := **m**) and intercellular interactions (**s** := **n**). E-G,J,K) Minimal stochastic pattern formation models, with black and white hexagons depicting final properties. E,F,J,K) start with random ***g***_0_, while G) starts with an imperfect pattern in which only a fraction of initial properties are scrambled. Top: Representative final patterns. Where shown, purple dots indicate fixed extracellular cue values *m*(***x***) and trajectories change from cold to warm colors over time. Bottom: time series plots of PI-generating information flows (solid curves, 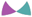) and upper bounds (dashed and dotted curves) associated with candidate information sources (Eq. (9)). E) A morphogen gradient **∇***m*(***x***) is sufficient to generate PI through instruction. F) A directional motion bias **s** := **d** derived from *m*(***x***) is sufficient to generate PI through sorting. G) Nearest-neighbor interactions refine a pattern. Black and white dots indicate ***g***_0_. Majority properties in local neighborhoods, **s** := **n**, are sufficient to generate PI through instruction. H) Programmed and self-organized patterning are distinguished by the extent to which information flows between cells to generate PI. I) The self-organizing mechanism in (G) amplifies *I*(**x**_*t*_; **g**_*t*_) *> I*(**x**_0_; **g**_0_) *>* 0. J) Combines mechanisms in (E) and (G) with a noisier *m*(***x***). Sources **m** and 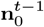 are jointly sufficient for PI generation. **m** alone is insufficient, as *I*(**x**; **m**|**g** ) does not bound *I*(**x**; **g**_*t*_|**g**_0_ ). K) Tissue geometry and differential adhesion facilitate self-organized PI generation through sorting. A cumulative directional swapping bias 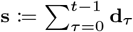 derived from adhesion differences in local neighborhoods is sufficient. See Section S3.4 for model details, and [Movie 1](https://www.dropbox.com/scl/fi/keokfrys8n6wull67esnx/all_mechanisms_movie_updated.mp4?rlkey=u3lwpvmja4tt86jwse9ff9798&st=dmoingv8&dl=0) for the time evolution of each model.

The final terms of Eqs. (8a-8b) (terms 5-6 of Eq. (2)) are forward-looking CMI terms that isolate directed information flow between **x** and **g**, distinguishing it from the mere preservation of initial correlations [21, 22] (Section S3.3). They are related to transfer entropy [21,66], but quantify finite-time information flow. After quantifying these flows, tracing PI generation to mechanisms requires identifying information sources (**s**) for which **x** or **g** serves as a proxy (Fig. 4C). The hypothesis that **s** is a sufficient information source for PI generation implies (Section S3.1):

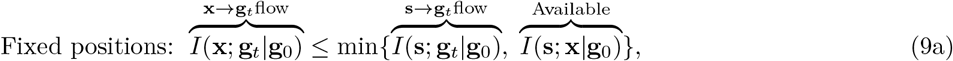

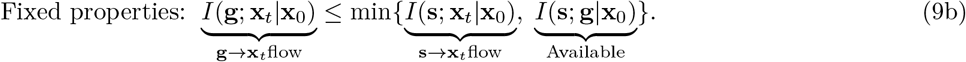

These conditions require that an information source both predict changes in one variable and provide new information about the other. If the corresponding inequality is violated, then **s** is insufficient to account for the generated PI, falsifying the hypothesis and requiring other information sources. In general, any PI-generating information sources must reside either in extracellular cues (**s** := **m**, e.g., morphogens) or in intercellular interactions (**s** := **n**, e.g., with neighboring cells) (Fig. 4D). Below, we model 2D hexagonal lattices of cells that update either **g**_*t*_ or **x**_*t*_ (subject to Eqs. (8–9)) via stochastic rules, illustrating how generated PI can be traced to information sources (Fig. 4E–K).

### 3.2 Information from extracellular cues

Extracellular information sources **s** := **m** canonically include morphogens and more generally include any molecular or mechanical cues external to the cellular pattern. These cues may instruct cell property changes or facilitate sorting by biasing cell motility. In Fig. 4E, patterns form under a noisy, fixed *m*(***x***). Eqs. (2) and (8a) identify PI generation via instruction, and Eq. (9a) verifies that **m** provides PI *I*(**m**; **x**|**g**_0_) that can flow to properties *I*(**m**; **g**_*t*_|**g**_0_) to generate PI. In Fig. 4F, cells move through *m*(***x***). Eqs. (2) and (8b) identify PI generation via sorting. Eq. (9b) verifies that a directional cell motion bias (**d**) derived from **∇***m*(***x***) can provide directional information [67] to each cell type *I*(**d**_0_; **g**|**x**_0_), and that exposure to this cue over time predicts final positions 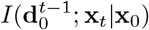, suggesting chemotaxis.

In Fig. 4F, cells sample cues while moving, making them dynamic in their co-moving frames [53, 68, 69]. When cues are dynamic, relevant information may reside not in instantaneous values, but in time series 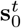 [7]. Generally, time series—joint distributions over multiple times—contain *temporal* features (e.g., frequency, duration) that are absent in instantaneous snapshots. Cells might perform various computations to extract relevant information [70, 71], as in the mouse neural tube, where 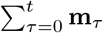 recovers most PI available in 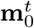, suggesting temporal integration as an efficient interpretation mechanism [7]. Feedback on **m** may be cell-autonomous, as in autocrine signaling, or depend on other cells, as in Turing patterns [72, 73] and self-generated gradients [74, 75]. Purely cell-autonomous feedback (Fig. 4H, top) amounts to local data processing, which cannot increase PI [19], whereas non-cell-autonomous feedback on **m** (Fig. 4H, bottom) can increase PI [76] by mediating intercellular interactions.

### 3.3 Information from intercellular interactions

Intercellular interactions require no external cues. Instead, cells’ properties depend on their neighbors’ properties [77–79] (Fig. 4D, H). We formalize this dependence by introducing an information source **s** := **n** ∼ *p*(***n***) for a cell’s nearest neighbors ***n*** ∈ 𝒩. In Fig. 4G, we consider a simple inductive mechanism in a static tissue in which cells adopt the majority state of their neighbors. When *I*(**x**_0_; **g**_0_) = 0, this mechanism fails to generate PI, but when *I*(**x**_0_; **g**_0_) *>* 0 it can increase PI (Fig. 4I) by correcting local pattern errors, analogous to a repetition code. Its generative capacity relies on *I*(**n**_0_; **x**|**g**_0_), enabling error identification, and *I*(**n**_0_; **g**_*t*_|**g**_0_), enabling correction. Sorting via intercellular interactions [80] can similarly correct patterning errors [81, 82]. When intercellular interactions complement extracellular cues, they may be individually inadequate but jointly sufficient. Fig. 4J combines the mechanisms in Fig. 4E,G with a noisier *m*(***x***). With both mechanisms present, *I*(**x**; **m**|**g**_0_) fails to bound the rising *I*(**x**; **g**_*t*_|**g**_0_), which is instead traceable to the joint source history 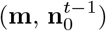. Tissue geometry [83, 84] can also provide an external source of information. In Fig. 4K, cells sort via differences in homotypic adhesion to generate a bullseye pattern [85–87]. A cumulative directional bias, 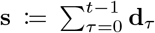, provides a sufficient information source (Fig. 4K, Section S3.4). This bias originates at the boundary, where lower homotypic adhesion is energetically favored, and spreads inwards, providing *I*(**s**; **g**|**x**_0_), predicting *I*(**s**; **x**_*t*_|**x**_0_), and gradually generating PI from *I*(**x**_0_; **g**_0_) = 0.

Without pre-existing PI or external information sources, intercellular interactions can break symmetries (e.g., A-P) at random [72], but cannot generate PI with respect to an established axis. Because development requires successive symmetry-breaking, embryos typically exhibit *mixed* self-organization [73], in which pre-existing PI provides a scaffold for self-organizing mechanisms to refine [11]. In Fig. 4G,I, the generated PI is traced to spatial correlations in **n**. In principle, **n** can extend beyond nearest neighbors to encode the joint properties and positions of all other cells, and thus information about the entire pattern. Patterns’ total correlation [3, 88]—capturing all their internal statistical dependencies—can therefore provide a rich information source to support PI generation [89]. However, only some of this information is relevant, and its accessibility depends on cells’ interaction ranges and computational repertoires.

## 4 Discussion

We have introduced a theoretical framework to quantify information flows in dynamic tissues from data and characterize how PI is preserved, lost, and generated over a finite time [0, *t*]—e.g., a developmental stage (Eq. (2)). These contributions are expressed as conditional mutual information terms among initial and final positions and properties (**x**_0_, **x**_*t*_, **g**_0_, **g**_*t*_)—capturing ubiquitous biological processes, including PI loss through positional mixing and property scrambling, and PI generation through instruction and sorting (Fig. 1).

Cell motion—central to PI dynamics—is becoming available at single-cell resolution in whole embryos. From our decomposition, we obtain information-theoretic measures of cell mixing that can be estimated directly from data. These include spatiotemporal mixing maps (Fig. 3A–C) and global predictive information *I*(**x**_0_; **x**_*t*_) (Eq. (5), Fig. 3D–E). When cell properties evolve slowly relative to—and do not affect—cell motion, dynamic tissues can be characterized as information channels in which an initial pattern must withstand cell mixing to preserve PI. In this setting, the final PI is upper-bounded not only by the initial PI but also by *I*(**x**_0_; **x**_*t*_) (Fig. 2C, Eq. (4)), constraining transmission of any pattern. In dynamic tissues, Lagrangian coordinates co-moving with the tissue provide a natural representation of position [43, 53] (Fig. 2D), consistent with the implicit coordinates used in fate maps. Using these coordinates, we find that morphogenesis—captured by the Cauchy–Green strain tensor [43]—structures regional mixing (Eq. (7)), enhancing it across multicellular attractors and suppressing it across repellers (Fig. 3F–H). This reveals an internal constraint [59, 60] between morphogenesis and PI dynamics.

When position and property dynamics are coupled, PI can be generated through information flows from initial positions to final properties **x**_0_ → **g**_*t*_ (instruction), initial properties to final positions **g**_0_ → **x**_*t*_ (sorting), and along trajectories (Eqs. (2,8)). We derive information-theoretic inequalities that test whether candidate information sources—extracellular cues or intercellular interactions—suffice to account for these PI-generating information flows (Eq. (9), Fig. 4A–D), quantitatively distinguishing programmed and self-organized strategies [11, 83]. Using minimal 2D tissue models, we illustrate how PI can be traced to extracellular information sources (e.g., morphogens), and how intercellular interactions and generic physical mechanisms such as differential adhesion [90] can generate PI from neighborhood correlations and tissue geometry (Fig. 4E-K). We note that our framework applies to any cell property: from molecular (e.g., gene expression) to mechanical [91], morphometric [92], or topological [93] observables.

By treating embryos as stochastic dynamical systems, our framework elucidates how the interplay between position and property dynamics drives pattern formation in dynamic tissues. This connects the kinematics and mechanics of cell motion to tissue patterning within a unified framework for embryo-scale information processing, naturally interpreted across Marr’s levels of analysis [94]. Increasing PI is a *computational* problem [11]; instructing and sorting—along with mitigating mixing and scrambling—encompass alternative *algorithms*; and tracing information flows to their sources constrains the space of sufficient mechanistic *implementations*. Understanding how information is processed and amplified through embryogenesis [95, 96] to pattern dynamic tissues remains a fundamental challenge, for which our framework offers a new theoretical foundation.

## Acknowledgments

We thank Miloš Nikolić and Enrico Maiorino for feedback and for helpful discussions. MS acknowledges support from NSF PHY-2413073, NIH R35GM156889, and HFSP Early Career Grant RGEC31/2024.

## Supplementary Materials

### S1 Supplement to Section 1

#### S1.1 Derivation of Eq. (2)

We derive Eq. (2) using repeated applications of the (redundancy-positive) definition of interaction information [20]

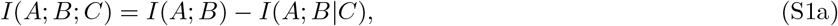

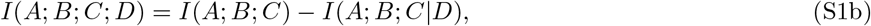

which follows from the chain rule for MI [19]. First, we use Eqs. (S1b)-(S1a) to expand the four-way interaction information *I*(**x**_0_; **g**_0_; **x**_*t*_; **g**_*t*_) in four ways that isolate either initial or final PI:

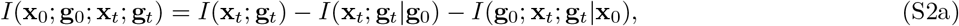

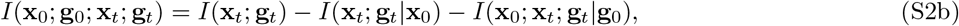

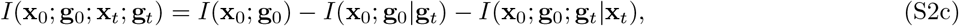

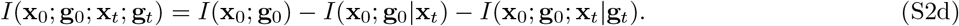

Averaging the two expansions involving final PI (Eqs. (S2a)-(S2b)) and rearranging, we obtain:

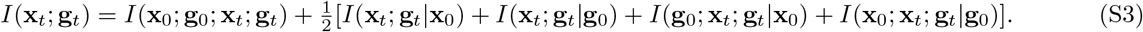

Conditioning all terms in Eq. (S1a) defines the conditional interaction information (CII), and rearranging yields:

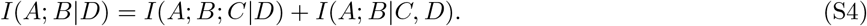

Applying Eq. (S4) to the CMI terms in Eq. (S3), we obtain:

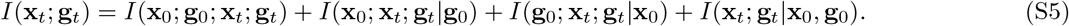

Averaging the two expansions involving initial PI (Eqs. (S2c)-(S2d)) and performing the same steps, we obtain:

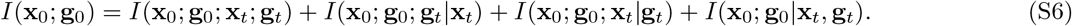

Subtracting Eq. (S6) from Eq. (S5) cancels *I*(**x**_0_; **g**_0_; **x**_*t*_; **g**_*t*_) and yields:

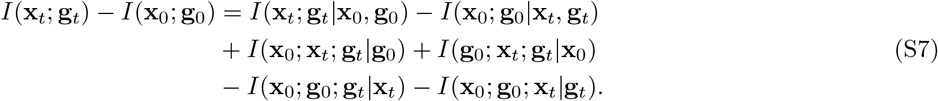

Eq. (S7) decomposes the change in PI into CMI terms for residual generation and joint loss, together with four CII terms corresponding to where only three of the four variables overlap in Fig. 1A. Eq. (S7) is appealing because each term directly shares in (initial or final) PI. However, while the CMI terms are strictly nonnegative, the CII terms may be positive or negative [20]. Expanding the four CII terms using Eq. (S4), we have:

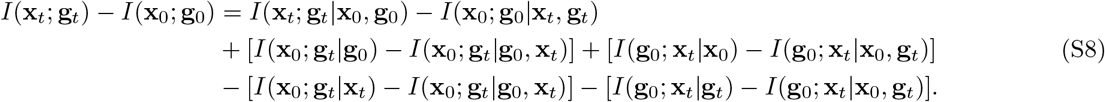

Finally, the *I*(**x**_0_; **g**_*t*_|**g**_0_, **x**_*t*_) and *I*(**g**_0_; **x**_*t*_|**x**_0_, **g**_*t*_) terms in Eq. (S8) cancel, yielding

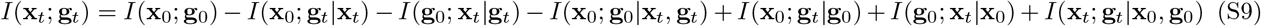

as desired. In Eq. (S9) (Eq. (2) in the main text), each term on the right-hand side is a CMI. Diagramatically, this decomposition, and each step in its derivation, is also consistent with any four-variable I-diagram 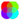.

#### S1.2 Minimal 1D patterning models

Each model in Fig. 1E-J was simulated with 1000 replicates on a 1D lattice with *N*_***x***_ = 20 positions, periodic boundaries, and *N*_*t*_ = 20 discrete time steps. Fig. S1 shows plots of each term in Eq. (2) over time for each patterning model. For Fig. 1E (preserving), positions and properties remained fixed over time. For Fig. 1F (mixing), properties remain fixed over time and, at each time step, a random set of disjoint neighbor pairs was selected and their positions were swapped. For Fig. 1G (scrambling), positions remain fixed over time and, at each time step, each cell independently flipped its property with probability *p* = 0.05. For Fig. 1H (instructing), positions remain fixed over time and, at each time step, cells independently updated their property with probability *p* = 0.3, setting it to *B* on the left half of the domain and to *W* on the right half. For Fig. 1I (sorting), properties remain fixed over time and, at each time step, all adjacent inversions (*W* immediately left of *B*) were identified, a random subset of non-overlapping pairs was selected, and each selected pair was swapped (*WB* → *BW* ). For Fig. 1J (correcting), cells mixed as in Fig. 1F and, at the end of each time step, cell properties were synchronously updated using elementary cellular automaton rule 232 [97].

**Fig. S1:**
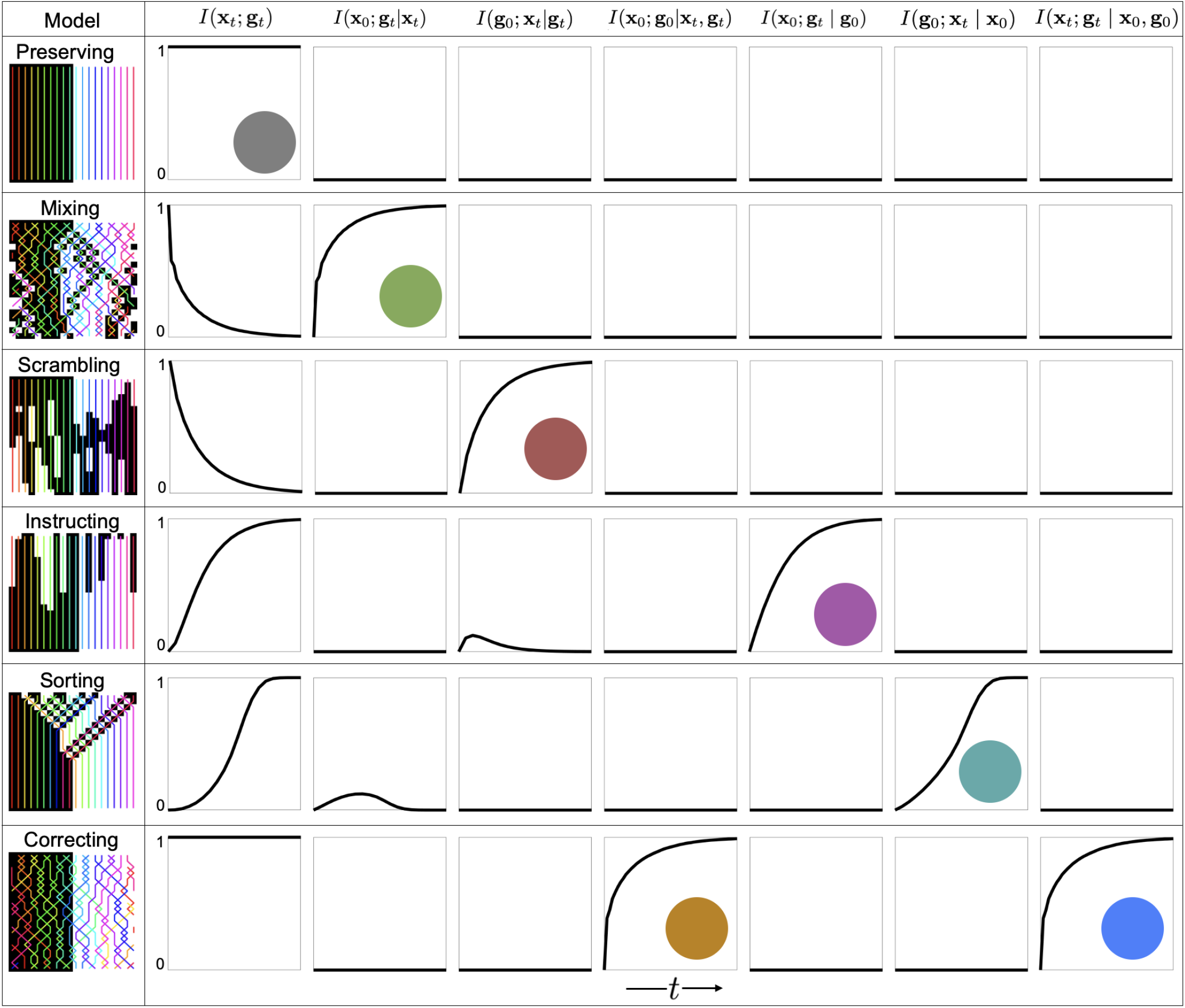
Positional information decomposition for minimal models. Time series of all time-dependent terms in Eq. (2) for each minimal model in Fig. 1E-J. Each row corresponds to one model, and each column to one term in the decomposition. Colored disks (colors as in Fig. 1) mark the relevant model in which each term reaches one bit.

#### S1.3 Algorithm for finite-time PI analysis

- Goal: estimate information-theoretic quantities from data on {**x**_0_, **x**_*t*_, **g**_0_, **g**_*t*_}.
- Given: data on positions and properties at times 0 and *t*, with cell identities linked through time.
- Discretize 𝒳 (positions) and 𝒢 (properties), trading off resolution against sample size.
  – 𝒳: discretize initial configuration. Infer 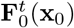 from i) average cell motion or ii) simultaneous measurement of continuously deforming, co-moving medium (e.g., ECM) to relate initial (**x**_0_) and final (**x**_*t*_) bins.
  – 𝒢: data may be continuous (e.g., gene expression measurements) or categorical (already discretized).
- Convert lineage data into distinct samples (cell trajectories). A cell must be present at *t* and traceable back to time 0—i.e., through a lineage—to count as a sample, providing a realization of {***x***_0_, ***x***_*t*_, ***g***_0_, ***g***_*t*_*}*. Initial cells that divide may contribute multiple samples. Those whose lineages begin after 0 or disappear before *t* are excluded.
- Let *N* be the total number of samples. Let 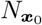 denote the number of samples in bin ***x***_0_. Similarly define 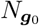, 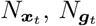, and joint counts 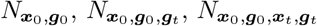, etc. (matrices and tensors).
- Estimate probabilities from frequencies: 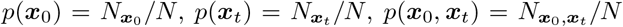 and likewise for other joint distributions. Conditional probabilities follow: *p*(***x***_*t*_|***x***_0_) = *p*(***x***_*t*_, ***x***_0_)*/p*(***x***_0_).
- Substituting these empirical probabilities into the corresponding information-theoretic expressions yields plug-in estimates. For example,

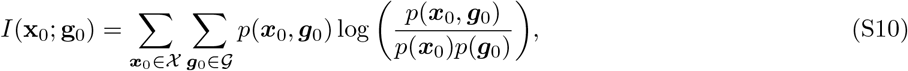

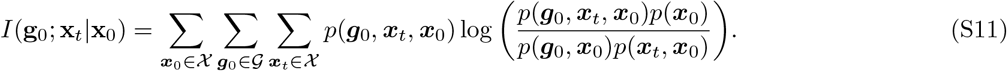

#### S1.4 PI loss and generation interpretations

Equation (2) is an exact decomposition for any joint distribution over {**x**_0_, **g**_0_, **x**_*t*_, **g**_*t*_*}*, and involves only nonnegative terms (MI and CMI). However, interpreting these terms as distinct bits of PI lost or generated depends on redundancy and synergy among the four variables, and is cleanest when synergy is absent or negligible. Synergy can appear in the interaction information between three or more variables. For example, the interaction information *I*(*A*; *B*; *C*) = − *I*(*A*; *B*) − *I*(*A*; *B*|*C*) (Eq. S1a) can be negative if conditioning on *C* increases the dependence between *A* and *B*, indicating synergy-dominated dependence, or positive, indicating redundancy-dominated dependence [20]. When synergistic interactions are present, the areas of information atoms for three or more variables in I-diagrams no longer provide a measure in bits [16]. For example, in Eq. (S7) the four CII terms may therefore become negative through large *I*(**x**_0_; **g**_*t*_|**g**_0_, **x**_*t*_) and *I*(**g**_0_; **x**_*t*_|**x**_0_, **g**_*t*_). The terms *I*(**x**_0_; **g**_*t*_|**g**_0_, **x**_*t*_) and *I*(**g**_0_; **x**_*t*_|**x**_0_, **g**_*t*_) cancel in Eq. (S8) and correspond to the two information atoms absent in the I-diagram in Fig. 1A. Both terms vanish when **x** or **g** is fixed, and also in the minimal models in Fig. 1E-J, implying that the CII terms are redundancy-dominated and individually reduce to non-negative CMIs. When nonzero, Eq. (S8) shows that they reallocate bits between PI loss (overlapping with **x**_0_ and **g**_0_) and generation (overlapping with **x**_*t*_ and **g**_*t*_) on account of differences in the 4th variable missing in each 3-variable CMI. In these cases, the 3-variable CMI terms cannot be interpreted as pure PI loss or pure PI generation. What always remains unambiguous are the differences *I*(**x**_0_; **g**_*t*_|**g**_0_) − *I*(**x**_0_; **g**_*t*_|**x**_*t*_) and *I*(**g**_0_; **x**_*t*_|**x**_0_) − *I*(**g**_0_; **x**_*t*_|**g**_*t*_), which are independent of the omitted atoms and quantify, respectively, the net position-to-property and property-to-position contributions to changing PI in Eq. (2).

### S2 Supplement to Section 2

#### S2.1 Isolating PI preservation

To isolate PI preservation, we must eliminate PI generation. Suppose **x** and **g** have independent dynamics, yielding two independent Markov processes **x**_0_ → **x**_*t*_ and **g**_0_ → **g**_*t*_ with no shared external drivers. Equivalently, we have a product channel [19] from (**x**_0_, **g**_0_) to (**x**_*t*_, **g**_*t*_): *p*(***x***_*t*_, ***g***_*t*_|***x***_0_, ***g***_0_) = *p*(***x***_*t*_|***x***_0_)*p*(***g***_*t*_|***g***_0_). Then, *I*(**x**_0_; **g**_*t*_|**g**_0_) = 0, *I*(**g**_0_; **x**_*t*_|**x**_0_) = 0, and *I*(**x**_*t*_; **g**_*t*_|**x**_0_, **g**_0_) = 0. Intuitively, **x**_0_ provides no information about **g**_*t*_ beyond **g**_0_, **g**_0_ provides no information about **x**_*t*_ beyond **x**_0_, and, conditioned on either initial variable, **x**_*t*_ and **g**_*t*_ are independent. Eq. (2) becomes

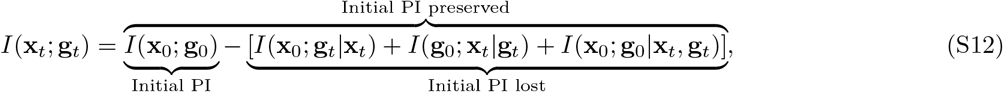

and because the CMI terms are nonnegative, *I*(**x**_*t*_; **g**_*t*_) ≤ *I*(**x**_0_; **g**_0_), clarifying that PI generation requires interactive position-property dynamics. We obtain alternative decompositions (Eq. (3)) via the chain rule for MI [19]:

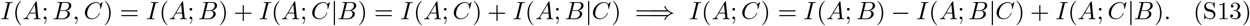

Applying Eq. (S13) to the two triplets (*A, B, C*) = (**x**_*t*_, **x**_0_, **g**_*t*_) and (**g**_*t*_, **g**_0_, **x**_*t*_) yields general identities, prior to imposing the independent-dynamics assumption:

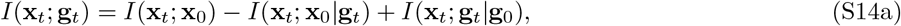

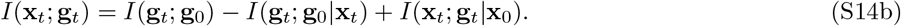

Because *p*(***x***_*t*_, ***g***_*t*_|***x***_0_, ***g***_0_) = *p*(***x***_*t*_|***x***_0_)*p*(***g***_*t*_|***g***_0_), the final terms of Eqs. (S14a)-(S14b) vanish: *I*(**x**_*t*_; **g**_*t*_|**g**_0_) = 0 and *I*(**x**_*t*_; **g**_*t*_|**x**_0_) = 0. This yields two simple alternatives to Eq. (S12) in terms of predictive information instead of initial PI:

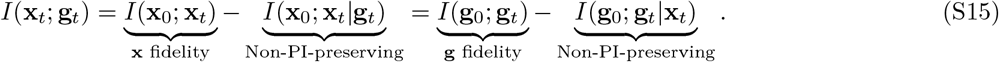

Eq. (S15) (Eq. (3) in the main text) reveals that each predictive-information term upper-bounds *I*(**x**_*t*_; **g**_*t*_), regardless of the initial pattern. Eq. (S12) and Eqs. (S15) hold for models in Fig. 1E-G because each satisfies our conditional independence assumptions. Under the product-channel assumption, all higher-order atoms in the corresponding I-diagrams (Fig. 2A–B) reflect inherited redundancy rather than new synergy, and so their areas are directly interpretable in bits.

#### S2.2 Channel capacities for positional information

Shannon showed that any communication channel has a capacity *C*, the maximum information that can be transmitted reliably under a specified noise model and fidelity criterion [12, 23]. In development, PI provides the fidelity criterion and mixing and scrambling arise from intrinsic and extrinsic process noise. Pattern maintenance over a developmental stage can therefore be conceptualized as transmission through a constant channel whose transition-probabilities are defined over the corresponding time interval. Because the embryo is a non-autonomous dynamical system [53], these transition probabilities vary over time, so distinct developmental stages define distinct channels. When developmental processes generate PI, they are no longer isolated channels, but their position or property dynamics may still be investigated as such to characterize the effects of noise for which PI generation mechanisms compensate. In Section 2, we consider channels with fixed properties and changing positions (Eq. (4)) where the input is an encoding *p*(***g***|***x***_0_) and the output is **x**_*t*_, from which the input can be inferred to the extent that PI is preserved. Conversely, for independently changing properties in a static tissue,

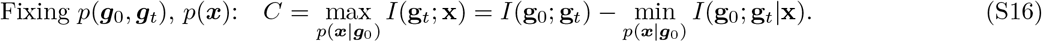

The predictive information *I*(**g**_0_; **g**_*t*_) quantifies the fidelity of property mappings along cell trajectories and lineages, limited, for example, by stochasticity in gene expression [98] dispersing *p*(***g***_*t*_|***g***_0_) within lineages, analogous to mixing in gene expression space. Finally, when neither properties nor positions are fixed but the dynamics define a product channel, both capacities still follow from Eq. (3), but the extrema are taken over *p*(***g***_*t*_|***x***_0_) and *p*(***x***_*t*_|***g***_0_) rather than over prepattern encodings. In all these cases, the ratio *I*(**x**_*t*_; **g**_*t*_)*/C* ≤ 1 provides a natural measure of the coding efficiency of observed prepatterns.

#### S2.3 Regional flow map estimation

We estimate a coarse flow map 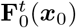 by advecting a regular 3D grid using cell trajectory data. First, we construct an initial grid (configuration) covering the bounding box of the trajectory data. The initial grid spacing is set to 10 *µ*m for Drosophila, 26 *µ*m for zebrafish, and 30 *µ*m for mouse to finely resolve the average tissue flows. At each time step *t*_*k*_ → *t*_*k*+1_, displacement samples are assembled from i) trajectories present at both *t*_*k*_ and *t*_*k*+1_ and ii) lineage births at *t*_*k*+1_, with a displacement from the mother’s position at *t*_*k*_ to the newly appearing daughter position at *t*_*k*+1_, ensuring that cell division contributes to the coarse flow. Grid vertex displacements are then estimated by Gaussian kernel averaging of nearby samples using a KD-tree neighborhood search. The kernel radius is initialized to the grid spacing and expanded adaptively until at least 12 displacement samples are included. Grid advection is performed with second-order Runge-Kutta (midpoint) integration, yielding a flow map for all grid vertices. This vertex map defines the flow map 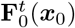 of the material regions defined by their corner vertices, initially cubes. Trajectory occupancy within these deforming regions is then used to estimate *p*(***x***_0_, ***x***_*t*_) (Section S1.3). Coarsened material regions—larger, non-overlapping cubes on the initial configuration—provide probability estimates across spatial resolutions.

#### S2.4 Information estimate validation

Mutual information is challenging to estimate accurately from finite samples. For Gaussian variables, or when the form of a distribution can be assumed, parametric mutual information estimators may be used. Here, however, we cannot assume a parametric form, nor do we have a continuous representation of the deformed domain, which itself must be constructed from finite samples. We therefore avoid estimators that rely on smooth underlying probability densities or metric structure in the deformed configuration. Instead, for all values reported in Fig. 3A-D,F we use the plug-in estimator, which computes probabilities directly from the empirical frequencies (Section S1.3). In Fig. 3A-F, each estimate is associated with a spatial resolution on the initial configuration, constraining pattern preservation at that resolution (Eq. (4)). The plug-in estimator exhibits an upward finite-sample bias for mutual information that increases with the number of bins and decreases with the number of samples (*N* cell trajectories) [99]. Because we require cell trajectories to be present in the data at both 0 (fixed initial time) and *t* (advancing final times), the sample size decreases as *t* increases, leading to larger estimation bias at later times. To leading orders in 1*/N*,

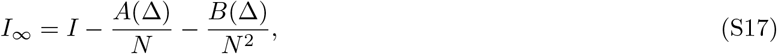

where *I*_∞_ is the infinite-sample limit and *A, B* depend on the bin resolution Δ. Following standard finite-sample extrapolation procedures [99–101], we compute *I*(*N* ) by averaging estimates from different-sized random subsets of the cell trajectories and fit Eq. (S17) to extrapolate *I*_∞_ as *N*→ ∞ . This procedure i) improves *I*(**x**_0_; **x**_*t*_) estimates for a given resolution and time interval, and ii) identifies resolutions and time intervals for which estimation is less reliable, indicated by nonlinearity or a poor fit of Eq. (S17). Fig. S2 shows the extrapolation behavior at five times across the spatial resolutions shown in Fig. 3D for *Drosophila*, mouse, and zebrafish gastrulation. We also perform the same procedure with randomly permuted ***x***_*t*_ labels—pairing random trajectory start and endpoints—to destroy any dependence in the data. In this shuffled null, fitting Eq. (S17) should extrapolate to *I*_∞_ (**x**_0_; **x**_*t*_) = 0. Extrapolation of shuffled data to a positive value suggests residual upward finite-sampling bias not attributable to a real signal. Fig. S2 demonstrates that, because *N* decreases with increasing *t*, mutual information estimates remain reliable only up to intermediate times in zebrafish and *Drosophila* gastrulation. Accordingly, in Fig. 3A-C, we plot prospective information values up to intermediate times in zebrafish (15 hpf) and *Drosophila* (4.2 *h* AEL) gastrulation. For visualization, and to give each embryo similar initial prospective information values, we plot at spatial resolutions of 69 *µm* for *Drosophila*, 107 *µm* for zebrafish, and 86 *µm* for mouse gastrulation. In Fig. 3E, we filter estimates by requiring that the shuffled extrapolation remains below 0.1 bits, excluding some later times and higher resolutions.

#### S2.5 Dynamic morphoskeletons

For a given time interval [0, *t*], the dynamic morphoskeleton [43]—consisting of attractors and repellers—is determined from the largest finite-time Lyapunov exponent (FTLE) in forward and backward time, defined as

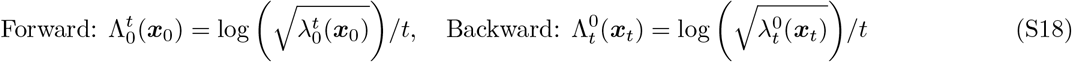

**Fig. S2:**
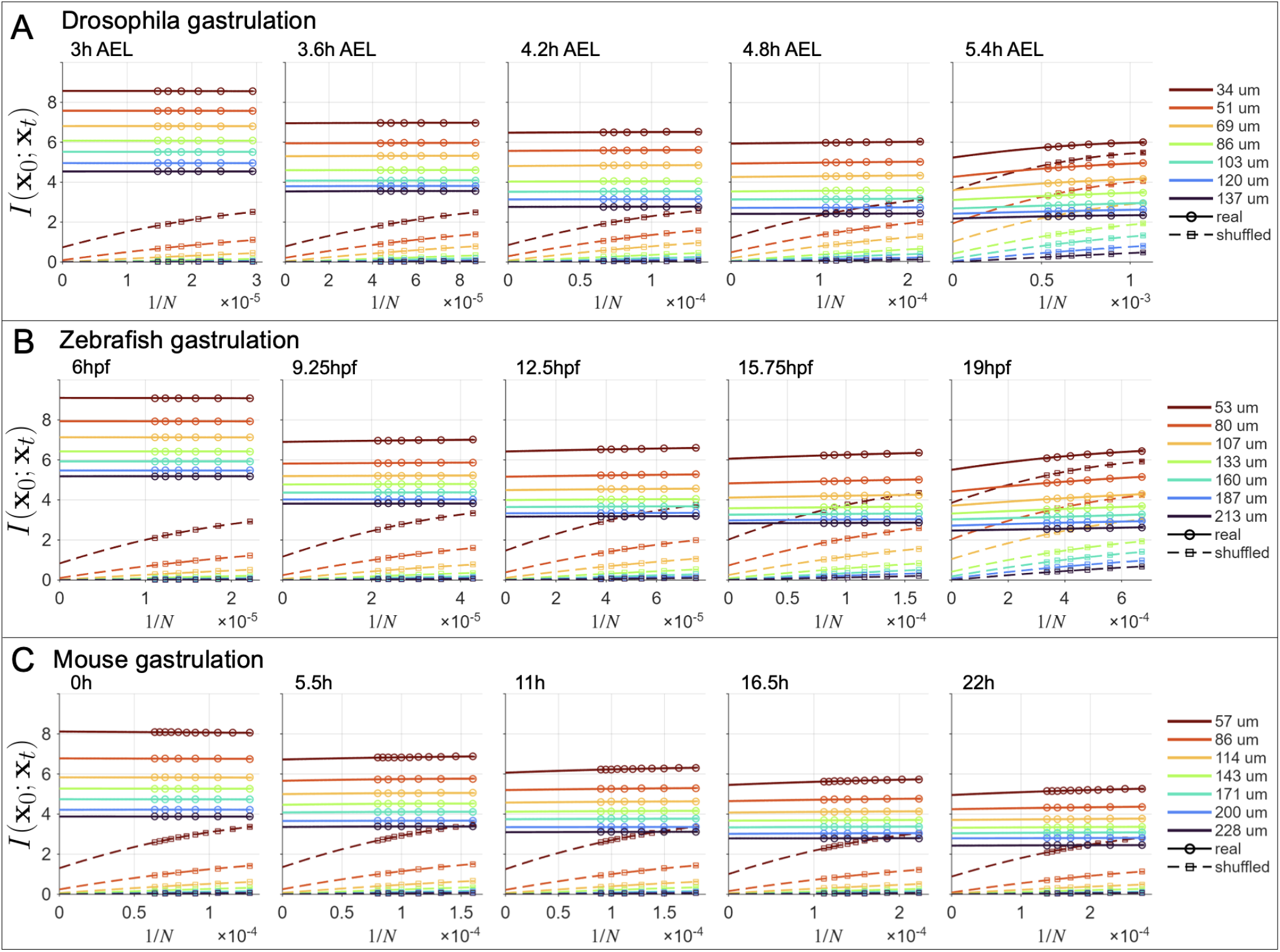
Predictive information estimation. Extrapolation tests for estimating *I*(**x**_0_; **x**_*t*_) at multiple spatial resolutions and times using the real cell trajectory data and shuffled-null data. Points show averages over 100 random subsets containing 50% − 100% of the samples. Solid and dashed lines show quadratic fits to the real and shuffled estimates. A) *Drosophila* gastrulation. B) Zebrafish gastrulation. C) Mouse gastrulation.

where 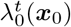 and 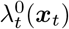 are the largest eigenvalues of 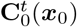 and 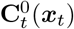, respectively. Plotting scalar fields 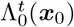 and 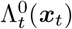 on the initial and final configurations reveals repellers and attractors identified by high FTLE values (Fig. 3F) [43]. Note that attractors and repellers are kinematic structures in the dynamical systems sense, and do not imply causative mechanisms such as chemo-attractants, only coherent structures in the tissue flows.

#### S2.6 Expanded form of Eq. (7)

For generality, we consider a compressible flow **v**(***x***, *t*) and heterogeneous, anisotropic, and time-dependent diffusivity **D**(***x***, *t*). We use Lagrangian coordinates ***x***_0_ induced by the regional flow map 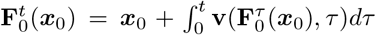. Recognizing that a conditional probability density 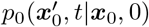 (hereafter *p*_0_) over the domain is mathematically equivalent, up to normalization, to a mass-conserved concentration field 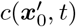, we have from [53, 56] that

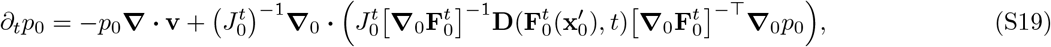

where 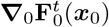 is the deformation gradient, i.e., the Jacobian of the flow map with respect to ***x***_0_, and 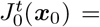 det 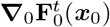 is the Jacobian determinant, a volume ratio *V*_*t*_*/V*_0_ that deviates from 1 only through compressibility. While probabilities are dimensionless, probability densities—used in Eqs. (6), (7), and (S19)—inherit the inverse units of their sample space. In a *d*-dimensional position space, they have units [*L*]^−*d*^. For incompressible motion, Lagrangian and flow-mapped Eulerian probability densities coincide. When 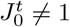, they use different differential volume elements, and conservation of probability requires 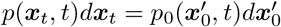. Since 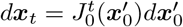, it follows that

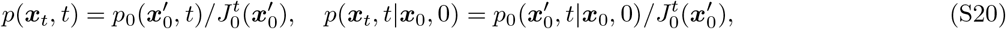

allowing conversion between probability densities over initial and final configurations for compressible flows. This coordinate transformation affects differential entropies, but the 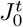 Jacobian terms cancel in relative entropies such as specific information and mutual information, which are invariant under smooth, invertible changes of variables [19].

#### S2.7 Predictive information dynamics

We illustrate the simple case of an incompressible dynamic tissue with uniform initial cell density and drift-diffusion dynamics (Eqs. (6)-(7)). Because the set of regions is fixed, the sample space 𝒳 is time-independent. Differentiating the prospective information for a specific initial region ***x***_0_ gives

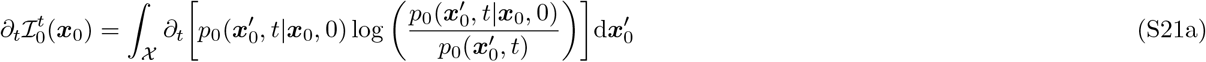

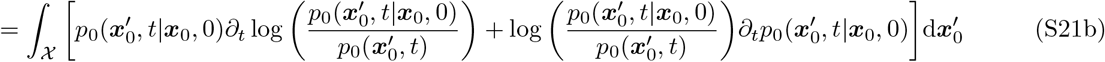

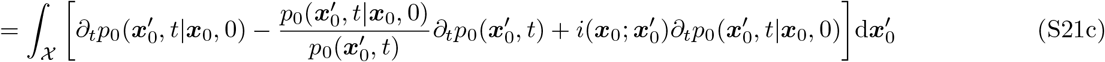

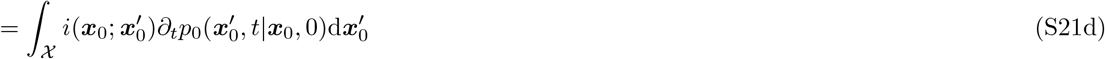

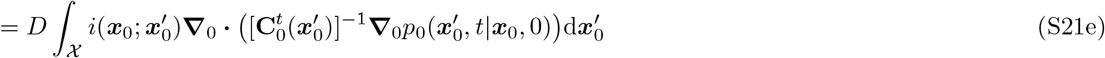

where we used normalization 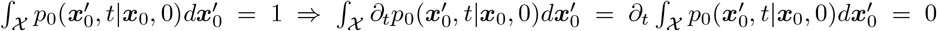 together with the fact that 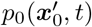 remains fixed, so 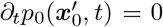. In the last line we substituted Eq. (7) with fixed 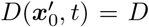. Relaxing the incompressibility and uniform initial density assumptions yields additional terms and requires substitution of the expanded form Eq. (S19) instead of Eq. (7). Eq. (S21e) pinpoints where prospective information decays faster or slower, depending on the interplay between random cell motility and 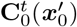. Taking the expectation of (S21e) over initial regions ***x***_0_ yields the finite-time predictive information dynamics

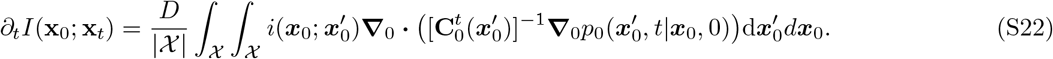

Taking the limit 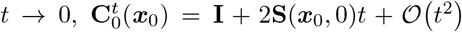 [102] in terms of the objective Eulerian rate of strain tensor 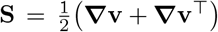, revealing that information transmission is affected only by frame-invariant features of the velocity field. Prior work demonstrated that rigid rotation does not affect entropy production [103] and that linear flows accelerate information loss [30, 104]. Eq. (S22) generalizes these observations to nonlinear flows, including time-dependent morphogenesis, and elucidates how deformation augments predictive information dynamics.

### S3 Supplement to Section 3

#### S3.1 Identifying sufficient information sources

In static tissues, information flow from **x** to **g**_*t*_ decomposes—via Eq. (S4)—with respect to an information source **s**:

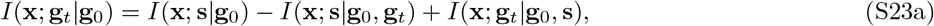

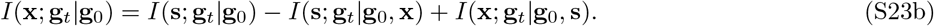

If **s** is a sufficient information source, then the residual term vanishes, *I*(**x**; **g**_*t*_|**g**_0_, **s**) = 0, and Eq. (9a) follows from the non-negativity of CMI. Likewise, with fixed cell properties, information flow from **g** to **x**_*t*_ decomposes as:

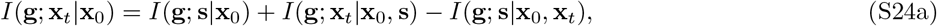

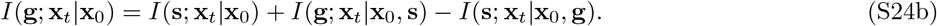

If **s** is sufficient, then *I*(**g**; **x**_*t*_|**x**_0_, **s**) = 0, and Eq. (9b) follows from the non-negativity of CMI. While one could test the sufficiency of **s** in each case by directly estimating *I*(**x**; **g**_*t*_|**g**_0_, **s**) and *I*(**g**; **x**_*t*_|**x**_0_, **s**), these residual terms each involve four variables, whereas the other terms involve only three, and are therefore easier to estimate from finite data.

#### S3.2 Information flows along trajectories

If both **x** and **g** change, PI generation also includes the residual term *I*(**x**_*t*_; **g**_*t*_|**x**_0_, **g**_0_) in Eq. (2). Two alternative algebraic decompositions—again via the chain rule for MI—absorb this residual term to assess information flows along trajectories, either from positional trajectories to final properties or property trajectories to final positions:

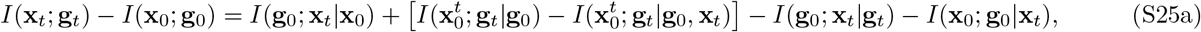

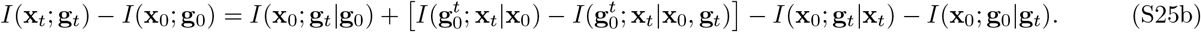

By the non-negativity of CMI, any increase in PI is upper-bounded—together with each original PI-generating information flow term from the initial variables in Eq. (2)—by information flows along trajectories 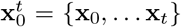 and 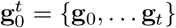, where additional information sources may be present:

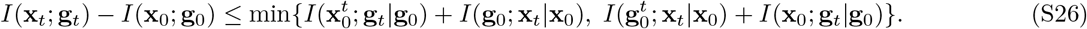

Although 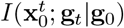 and 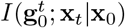 are not strictly past-to-future transfers—because 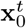 and 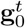 contain **x**_*t*_ and **g**_*t*_—they can be temporally decomposed into a transfer from an earlier trajectory segment and a residual term using the chain rule for MI,

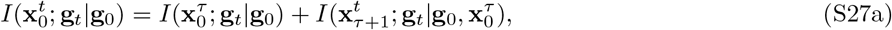

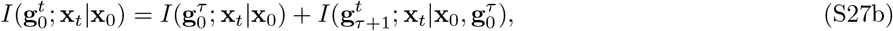

quantifying how different trajectory segments [0, *τ* ] ⊂ [0, *t*] contribute to PI generation. Residuals 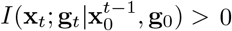 or 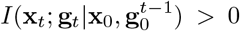 may indicate inadequate time resolution. Finally, decomposing these trajectory-based information flows with respect to a candidate source history 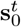 (e.g., extracellular cue exposure along trajectories):

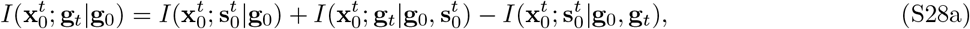

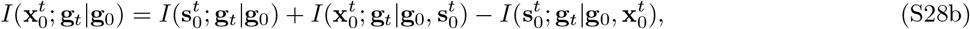

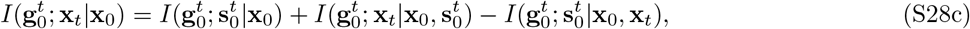

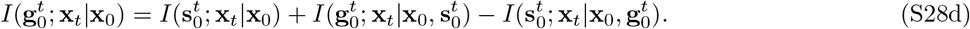

If 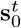 is a sufficient information source to predict property changes from spatial trajectories, then 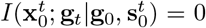 (instructing along trajectories). Likewise, if 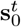 is a sufficient information source to predict positional changes from property trajectories, 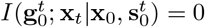 (sorting along trajectories). In this general case, the bounds become:

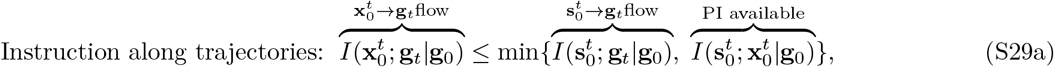

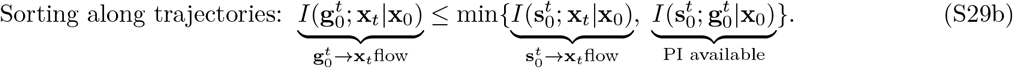

#### S3.3 Causal inference

Information-theoretic approaches to causal inference often assume long, stationary, ergodic time series to estimate transition probabilities. Development is instead governed by time-varying dynamical rules [105], yielding non-stationary time series. However, as in Section 2, coarse-graining can trade resolution for sample size within a single embryo, and reproducible, stage-specific dynamics can more generally permit stage-specific estimation from developmental ensembles insofar as they provide independent time-series replicates [106]. Comprehensively dissecting PI dynamics requires datasets of sufficient quantity, quality, and spatiotemporal resolution to estimate the relevant information-theoretic quantities. CMI supports limited observational causal inference, providing a model-free, nonlinear generalization of Granger causality [107, 108], but is not sufficient to establish mechanism. Multiple variables may contribute [20] and provide sufficient information. Because CMI alone does not distinguish among unique, redundant, and synergistic contributions from multiple information source, it may misattribute, underestimate, or overestimate causal influence [109]. Accordingly, Eqs. (9a)-(9b) can rule out candidate mechanisms acting in isolation, but cannot verify them, so this observational approach should be complemented by mechanistic modeling and, where possible, interventionist experimental perturbations. Conditioning on additional variables can further isolate information flows at the cost of increasing dimensionality and data requirements [110]. Conversely, compressing information sources—for example, by estimating information from scalar functionals of time-series [7], as in Section 3—can test more specific mechanistic hypotheses without incurring the full data demands of high-dimensional trajectory distributions. Explicit separation of unique, redundant, and synergistic contributions from multiple information sources—and fully disentangling PI loss and generation in the presence of synergistic interactions (Section S1.3)—would likely require partial information decomposition (PID) [111], which remains an area of active research [112].

#### S3.4 Two-dimensional models

Each model in Fig. 4E-G,J,K was simulated in 1000 independent replicates on a hexagonal lattice with *N*_***x***_ = 238 positions inside an approximately circular, non-periodic domain. Simulations for Fig. 4E-J used *N*_*t*_ = 20 time steps while simulations for Fig. 4K used *N*_*t*_ = 1000. For Fig. 4E-F, an external cue *m*(***x***) is generated independently for each replicate as a static exponential gradient 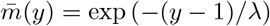 along the vertical (*y*) axis, with multiplicative lognormal noise 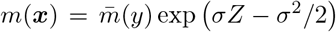 where *Z* ∼ 𝒩 (0, 1), i.i.d. across lattice sites. The gradient length scale is chosen so that 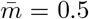 halfway across the domain. In Fig. 4E, *σ* = 0.1, and each cell updates its state with probability 0.3, becoming black if *m <* 0.5 and white otherwise. In Fig. 4F, *σ* = 0.02, and cells exchange positions via cue-biased nearest-neighbor swaps. At each time step, 10 sets of random non-overlapping nearest-neighbor pairings are executed sequentially. Each heterotypic pair swaps with probability 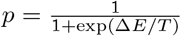 where Δ*E* = (*m*_2_ − *m*_1_)(*g*_2_ − *g*_1_) and *T* = 0.02, favoring swaps that move black cells toward higher *m* and white cells toward lower *m*. As a candidate information source, we defined the local directional bias **d**(***x, g***) ≡ (2***g*** − 1)∑_*j*_ (*m*(***x***_*j*_) − *m*(***x***))Δ*y*_*j*_, where Δ*y*_*j*_ is the vertical displacement to nearest neighbor *j*. For the information-theoretic analysis, we coarse-grained **x**_0_ and **x**_*t*_ into two vertical regions with approximately equal numbers of cells, and **d** into five bins with edges (−∞, −1, − 0.3, 0.3, 1, ∞ ) to estimate history variables 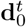. For Fig. 4G-H, a perfect pattern with *I*(**x**; **g**) = 1 is perturbed by randomizing a fraction of cell states, and cells then update with probability *p* = 0.3 to match the majority of their neighbors, with no change in the event of a tie. In Fig. 4G, 30% of cells are randomized initially. In Fig. 4H, we randomize with 100 uniformly spaced values of 0% −100% and 1000 replicates per value. For Fig. 4J, we combine the instructive gradient mechanism of Fig. 4E (*σ* = 0.12, update probability 0.4) and the neighbor majority mechanism of Fig. 4G (update probability 0.3) sequentially at each time step, coarse-graining **x** into two vertical regions for analysis. For Fig. 4K, local swap probabilities are computed from the current configuration *g*(***x***) based on homotypic and heterotypic neighbor contacts. The local adhesion energy is *E*_*T*_ (***x***) = *N*_0_(***x***)*E*_0_ + *N*_1_(***x***)*E*_1_ + *N*_≠_ (***x***)*E*_≠_, where *N*_0_(***x***), *N*_1_(***x***), and *N*_≠_ (***x***) are the numbers of homotypic black, homotypic white, and heterotypic neighbors, and *E*_0_, *E*_1_, and *E*_≠_ are the corresponding energies. For neighbor direction *j*, we define 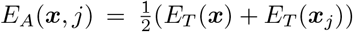 and *k*(***x***, *j*) = *k*_0_ exp (−*E*_*A*_(***x***, *j*)) with *k*_0_ = 15. At each time step, random non-overlapping nearest-neighbor pairs swap with probability *p*(***x***, *j*) = 1 − *e*^−*k*(***x***,*j*)^, as in Kawasaki dynamics [80,113]. As a candidate information source, we define the inward-outward swapping bias **d**(***x***) = **d**_out_(***x***) − **d**_in_(***x***), where **d**_out_ and **d**_in_ are the best local energy decreases achievable by outward and inward swaps. For analysis, we coarse-grain **x** into two radial regions and discretize **d** into five signed bins, accumulated along trajectories as 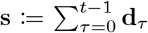. Here, boundary geometry initially biases the directionality of energetically favorable swaps for black and white cells at or near the boundary, accounting for the characteristic two-stage sorting process [86]: rapid boundary-driven rearrangements followed by slower bulk sorting.

## Movies

[Movie 1](https://www.dropbox.com/scl/fi/keokfrys8n6wull67esnx/all_mechanisms_movie_updated.mp4?rlkey=u3lwpvmja4tt86jwse9ff9798&st=dmoingv8&dl=0): Time evolution for realizations of the processes in Fig. 4E-G,J,K.

## References

[1] L. Wolpert, “Positional information and the spatial pattern of cellular differentiation,” Journal of theoretical biology, vol. 25, no. 1, pp. 1–47, 1969.

[2] J. O. Dubuis, G. Tkačik, E. F. Wieschaus, T. Gregor, and W. Bialek, “Positional information, in bits,” Proceedings of the National Academy of Sciences, vol. 110, no. 41, pp. 16301–16308, 2013.

[3] D. B. Brückner and G. Tkačik, “Information content and optimization of self-organized developmental systems,” Proceedings of the National Academy of Sciences, vol. 121, no. 23, p. e2322326121, 2024.

[4] L. McGough, H. Casademunt, M. Nikolić, Z. Aridor, M. D. Petkova, T. Gregor, and W. Bialek, “Finding the last bits of positional information,” PRX life, vol. 2, no. 1, p. 013016, 2024.

[5] M. Nikolić, V. Antonetti, F. Liu, G. Muhaxheri, M. D. Petkova, M. Scheeler, E. M. Smith, W. Bialek, and T. Gregor, “Scale invariance in early embryonic development,” Proceedings of the National Academy of Sciences, vol. 121, no. 46, p. e2403265121, 2024.

[6] A. Huang, L. Cocconi, B. Nicholls-Mindlin, C. Alexandre, G. Salbreux, and J.-P. Vincent, “A genetic circuit that extends the useful range of a bmp morphogen arose alongside insect wing evolution,” Current Biology, 2025.

[7] A. Marković, J. Briscoe, and K. M. Page, “Dynamics of positional information in the vertebrate neural tube,” Journal of the Royal Society Interface, vol. 21, no. 221, p. 20240414, 2024.

[8] T. Fulton, B. Verd, and B. Steventon, “The unappreciated generative role of cell movements in pattern formation,” Royal Society Open Science, vol. 9, no. 4, p. 211293, 2022.

[9] D. Pinheiro, R. Kardos, É. Hannezo, and C.-P. Heisenberg, “Morphogen gradient orchestrates pattern-preserving tissue morphogenesis via motility-driven unjamming,” Nature Physics, vol. 18, no. 12, pp. 1482–1493, 2022.

[10] T. Y.-C. Tsai and D. Pinheiro, “Coping with uncertainty: Challenges for robust pattern formation in dynamical tissues,” in Seminars in Cell & Developmental Biology, vol. 173, p. 103629, Elsevier, 2025.

[11] D. B. Brückner and G. Tkačik, “Marr’s three levels for embryonic development: information, dynamical systems, gene networks,” PRX Life, vol. 4, no. 1, p. 017001, 2026.

[12] C. E. Shannon, “A mathematical theory of communication,” The Bell system technical journal, vol. 27, no. 3, pp. 379–423, 1948.

[13] A. Levchenko and I. Nemenman, “Cellular noise and information transmission,” Current opinion in biotechnology, vol. 28, pp. 156–164, 2014.

[14] G. Tkačik and W. Bialek, “Information processing in living systems,” Annual Review of Condensed Matter Physics, vol. 7, no. 1, pp. 89–117, 2016.

[15] G. Tkačik and T. Gregor, “The many bits of positional information,” Development, vol. 148, no. 2, p. dev176065, 2021.

[16] R. W. Yeung, “A new outlook on shannon’s information measures,” IEEE transactions on information theory, vol. 37, no. 3, pp. 466–474, 1991.

[17] J. D. Farmer, “Information dimension and the probabilistic structure of chaos,” Zeitschrift für Naturforschung A, vol. 37, no. 11, pp. 1304–1326, 1982.

[18] W. Bialek, I. Nemenman, and N. Tishby, “Predictability, complexity, and learning,” Neural computation, vol. 13, no. 11, pp. 2409–2463, 2001.

[19] T. M. Cover, J. A. Thomas, et al., “Entropy, relative entropy and mutual information,” Elements of information theory, vol. 2, no. 1, pp. 12–13, 1991.

[20] W. McGill, “Multivariate information transmission,” Transactions of the IRE Professional Group on Information Theory, vol. 4, no. 4, pp. 93–111, 1954.

[21] T. Schreiber, “Measuring information transfer,” Physical review letters, vol. 85, no. 2, p. 461, 2000.

[22] K. Hlaváčková-Schindler, M. Paluš, M. Vejmelka, and J. Bhattacharya, “Causality detection based on information-theoretic approaches in time series analysis,” Physics Reports, vol. 441, no. 1, pp. 1–46, 2007.

[23] R. M. Fano and W. Wintringham, “Transmission of information,” 1961.

[24] O. Witteveen, S. J. Rosen, R. S. Lach, M. Z. Wilson, and M. Bauer, “Optimizing information transmission in the canonical wnt pathway,” arXiv preprint arXiv:2506.22633, 2025.

[25] G. Tkačik, C. G. Callan Jr, and W. Bialek, “Information capacity of genetic regulatory elements,” Physical Review E—Statistical, Nonlinear, and Soft Matter Physics, vol. 78, no. 1, p. 011910, 2008.

[26] M. Saez, G. Minas, E. Camacho-Aguilar, and D. A. Rand, “Measuring developmental information encoded by a dynamical landscape,” bioRxiv, 2026.

[27] T. A. Huijben, A. G. Anderson III, A. Sweet, E. Hoops, C. Larsen, K. Awayan, J. Bragantini, M. Lange, C.-L. Chiu, and L. A. Royer, “intracktive: a web-based tool for interactive cell tracking visualization,” Nature Methods, pp. 1–3, 2025.

[28] E. H. Linfoot, “An informational measure of correlation,” Information and control, vol. 1, no. 1, pp. 85–89, 1957.

[29] Y. Wan, J. El Kholtei, I. Jenie, M. Colomer-Rosell, J. Liu, Q. Zhang, J. N. Acedo, L. Y. Du, M. Codina-Tobias, M. Wang, et al., “Whole-embryo spatial transcriptomics at subcellular resolution from gastrulation to organogenesis,” Science, vol. 391, no. 6790, p. eadt3439, 2026.

[30] Y. Shi, R. Golestanian, and A. Vilfan, “Mutual information as a measure of mixing efficiency in viscous fluids,” Physical Review Research, vol. 6, no. 2, p. L022050, 2024.

[31] J. M. W. Slack, From egg to embryo: regional specification in early development. Cambridge University Press, 1991.

[32] M. Delarue, S. Sanchez, K. E. Johnson, T. Darribère, and J.-C. Boucaut, “A fate map of superficial and deep circumblastoporal cells in the early gastrula of pleurodeles waltl,” Development, vol. 114, no. 1, pp. 135–146, 1992.

[33] Y. Hatada and C. D. Stern, “A fate map of the epiblast of the early chick embryo,” Development, vol. 120, no. 10, pp. 2879–2889, 1994.

[34] P. J. Keller, A. D. Schmidt, J. Wittbrodt, and E. H. Stelzer, “Reconstruction of zebrafish early embryonic development by scanned light sheet microscopy,” science, vol. 322, no. 5904, pp. 1065–1069, 2008.

[35] F. Amat, W. Lemon, D. P. Mossing, K. McDole, Y. Wan, K. Branson, E. W. Myers, and P. J. Keller, “Fast, accurate reconstruction of cell lineages from large-scale fluorescence microscopy data,” Nature methods, vol. 11, no. 9, pp. 951–958, 2014.

[36] M. Lange, A. Granados, S. VijayKumar, J. Bragantini, S. Ancheta, Y.-J. Kim, S. Santhosh, M. Borja, H. Kobayashi, E. McGeever, et al., “A multimodal zebrafish developmental atlas reveals the state-transition dynamics of late-vertebrate pluripotent axial progenitors,” Cell, vol. 187, no. 23, pp. 6742–6759, 2024.

[37] J. Bragantini, I. Theodoro, X. Zhao, T. A. Huijben, E. Hirata-Miyasaki, S. VijayKumar, A. Balasubramanian, T. Lao, R. Agrawal, S. Xiao, et al., “Ultrack: pushing the limits of cell tracking across biological scales,” Nature Methods, pp. 1–14, 2025.

[38] K. McDole, L. Guignard, F. Amat, A. Berger, G. Malandain, L. A. Royer, S. C. Turaga, K. Branson, and P. J. Keller, “In toto imaging and reconstruction of post-implantation mouse development at the single-cell level,” Cell, vol. 175, no. 3, pp. 859–876, 2018.

[39] B. Bénazéraf, P. Francois, R. E. Baker, N. Denans, C. D. Little, and O. Pourquié, “A random cell motility gradient downstream of fgf controls elongation of an amniote embryo,” Nature, vol. 466, no. 7303, pp. 248–252, 2010.

[40] F. Xiong, W. Ma, B. Bénazéraf, L. Mahadevan, and O. Pourquié, “Mechanical coupling coordinates the co-elongation of axial and paraxial tissues in avian embryos,” Developmental Cell, vol. 55, no. 3, pp. 354–366, 2020.

[41] K. Uriu, R. Bhavna, A. C. Oates, and L. G. Morelli, “A framework for quantification and physical modeling of cell mixing applied to oscillator synchronization in vertebrate somitogenesis,” Biology open, vol. 6, no. 8, pp. 1235–1244, 2017.

[42] A. Mongera, P. Rowghanian, H. J. Gustafson, E. Shelton, D. A. Kealhofer, E. K. Carn, F. Serwane, A. A. Lucio, J. Giammona, and O. Campàs, “A fluid-to-solid jamming transition underlies vertebrate body axis elongation,” Nature, vol. 561, no. 7723, pp. 401–405, 2018.

[43] M. Serra, S. Streichan, M. Chuai, C. J. Weijer, and L. Mahadevan, “Dynamic morphoskeletons in development,” Proceedings of the National Academy of Sciences, vol. 117, no. 21, pp. 11444–11449, 2020.

[44] K. D. Irvine and E. Wieschaus, “Cell intercalation during drosophila germband extension and its regulation by pair-rule segmentation genes,” Development, vol. 120, no. 4, pp. 827–841, 1994.

[45] T. Stern, S. Y. Shvartsman, and E. F. Wieschaus, “Deconstructing gastrulation at single-cell resolution,” Current Biology, vol. 32, no. 8, pp. 1861–1868, 2022.

[46] A. McMahon, W. Supatto, S. E. Fraser, and A. Stathopoulos, “Dynamic analyses of drosophila gastrulation provide insights into collective cell migration,” Science, vol. 322, no. 5907, pp. 1546–1550, 2008.

[47] A. Francou, K. V. Anderson, and A.-K. Hadjantonakis, “A ratchet-like apical constriction drives cell ingression during the mouse gastrulation emt,” Elife, vol. 12, p. e84019, 2023.

[48] M. H. Dominguez, A. L. Krup, J. M. Muncie, and B. G. Bruneau, “Graded mesoderm assembly governs cell fate and morphogenesis of the early mammalian heart,” Cell, vol. 186, no. 3, pp. 479–496, 2023.

[49] J.-A. Montero, L. Carvalho, M. Wilsch-Brauninger, B. Kilian, C. Mustafa, and C.-P. Heisenberg, “Shield formation at the onset of zebrafish gastrulation,” 2005.

[50] F. Fagotto, “The cellular basis of tissue separation,” Development, vol. 141, no. 17, pp. 3303–3318, 2014.

[51] B. Monier, A. Pélissier-Monier, A. H. Brand, and B. Sanson, “An actomyosin-based barrier inhibits cell mixing at compartmental boundaries in drosophila embryos,” Nature cell biology, vol. 12, no. 1, pp. 60–65, 2010.

[52] D. Umetsu, B. Aigouy, M. Aliee, L. Sui, S. Eaton, F. Jülicher, and C. Dahmann, “Local increases in mechanical tension shape compartment boundaries by biasing cell intercalations,” Current Biology, vol. 24, no. 15, pp. 1798–1805, 2014.

[53] A. M. Plum and M. Serra, “Morphogen patterning in dynamic tissues,” PRX Life, vol. 3, no. 4, p. 043009, 2025.

[54] J. Ranft, M. Basan, J. Elgeti, J.-F. Joanny, J. Prost, and F. Jülicher, “Fluidization of tissues by cell division and apoptosis,” Proceedings of the National Academy of Sciences, vol. 107, no. 49, pp. 20863–20868, 2010.

[55] H. Risken, “Fokker-planck equation,” in The Fokker-Planck equation: methods of solution and applications, pp. 63–95, Springer, 1989.

[56] G. Haller, D. Karrasch, and F. Kogelbauer, “Material barriers to diffusive and stochastic transport,” Proceedings of the National Academy of Sciences, vol. 115, no. 37, pp. 9074–9079, 2018.

[57] C. Guillot, Y. Djeffal, M. Serra, and O. Pourquie, “Control of epiblast cell fate by mechanical cues,” bioRxiv, pp. 2024–06, 2024.

[58] H. Hamada, C. Meno, D. Watanabe, and Y. Saijoh, “Establishment of vertebrate left–right asymmetry,” Nature Reviews Genetics, vol. 3, no. 2, pp. 103–113, 2002.

[59] P. Alberch, “The logic of monsters: evidence for internal constraint in development and evolution,” Geobios, vol. 22, pp. 21–57, 1989.

[60] D. Cao, S. Garai, J. DiFrisco, and J. V. Veenvliet, “The logic of monsters: development and morphological diversity in stem-cell-based embryo models,” Interface focus, vol. 14, no. 5, p. 20240023, 2024.

[61] G. Struhl, K. Struhl, and P. M. Macdonald, “The gradient morphogen bicoid is a concentration-dependent transcriptional activator,” Cell, vol. 57, no. 7, pp. 1259–1273, 1989.

[62] T. Gregor, D. W. Tank, E. F. Wieschaus, and W. Bialek, “Probing the limits to positional information,” Cell, vol. 130, no. 1, pp. 153–164, 2007.

[63] H. M. McNamara, S. C. Solley, B. Adamson, M. M. Chan, and J. E. Toettcher, “Recording morphogen signals reveals mechanisms underlying gastruloid symmetry breaking,” Nature cell biology, vol. 26, no. 11, pp. 1832–1844, 2024.

[64] J. Pineau, J. Wong-Ng, A. Mayran, L. Lopez-Delisle, P. Osteil, A. Shoushtarizadeh, D. Duboule, S. Gobaa, and T. Gregor, “Fine-tuning mechanical constraints reveals uncoupled patterning and gene expression programs in murine gastruloids,” Development, vol. 152, no. 18, p. dev204711, 2025.

[65] J.-L. Maître, H. Turlier, R. Illukkumbura, B. Eismann, R. Niwayama, F. Nédélec, and T. Hiiragi, “Asymmetric division of contractile domains couples cell positioning and fate specification,” Nature, vol. 536, no. 7616, pp. 344–348, 2016.

[66] A. Tripathi, J. Dunkel, and D. J. Skinner, “Collective is different: Information exchange and speed-accuracy trade-offs in self-organized patterning,” PRX Life, vol. 4, no. 1, p. 013015, 2026.

[67] D. Fuller, W. Chen, M. Adler, A. Groisman, H. Levine, W.-J. Rappel, and W. F. Loomis, “External and internal constraints on eukaryotic chemotaxis,” Proceedings of the National Academy of Sciences, vol. 107, no. 21, pp. 9656–9659, 2010.

[68] T. Fulton, K. Spiess, L. Thomson, Y. Wang, B. Clark, S. Hwang, B. Paige, B. Verd, and B. Steventon, “Cell rearrangement generates pattern emergence as a function of temporal morphogen exposure,” 2021.

[69] K. Spiess, S. E. Taylor, T. Fulton, K. Toh, D. Saunders, S. Hwang, Y. Wang, B. Paige, B. Steventon, and B. Verd, “Approximated gene expression trajectories for gene regulatory network inference on cell tracks,” iScience, vol. 27, no. 9, p. 110840, 2024.

[70] M. Bauer, M. D. Petkova, T. Gregor, E. F. Wieschaus, and W. Bialek, “Trading bits in the readout from a genetic network,” Proceedings of the National Academy of Sciences, vol. 118, no. 46, p. e2109011118, 2021.

[71] M. Bauer and W. Bialek, “Information bottleneck in molecular sensing,” PRX Life, vol. 1, no. 2, p. 023005, 2023.

[72] A. M. Turing, “The chemical basis of morphogenesis,” Philosophical Transactions of the Royal Society of London. Series B, Biological Sciences, vol. 237, no. 641, pp. 37–72, 1952.

[73] J. Green and J. Sharpe, “Positional information and reaction-diffusion: two big ideas in developmental biology combine,” Development, vol. 142, no. 7, pp. 1203–1211, 2015.

[74] R. H. Insall, P. Paschke, and L. Tweedy, “Steering yourself by the bootstraps: how cells create their own gradients for chemotaxis,” Trends in Cell Biology, vol. 32, no. 7, pp. 585–596, 2022.

[75] J. Stock, T. Kazmar, F. Schlumm, E. Hannezo, and A. Pauli, “A self-generated toddler gradient guides mesodermal cell migration,” Science Advances, vol. 8, no. 37, p. eadd2488, 2022.

[76] P. Hillenbrand, U. Gerland, and G. Tkačik, “Beyond the french flag model: exploiting spatial and gene regulatory interactions for positional information,” PLoS One, vol. 11, no. 9, p. e0163628, 2016.

[77] I. Salazar-Ciudad, J. Jernvall, and S. A. Newman, “Mechanisms of pattern formation in development and evolution,” 2003.

[78] Y. Akieda, S. Ogamino, H. Furuie, S. Ishitani, R. Akiyoshi, J. Nogami, T. Masuda, N. Shimizu, Y. Ohkawa, and T. Ishitani, “Cell competition corrects noisy wnt morphogen gradients to achieve robust patterning in the zebrafish embryo,” Nature communications, vol. 10, no. 1, p. 4710, 2019.

[79] T. Dullweber and A. Erzberger, “Mechanochemical feedback loops in contact-dependent fate patterning,” Current Opinion in Systems Biology, vol. 32, p. 100445, 2023.

[80] R. M. Garner, S. E. McGeary, A. M. Klein, and S. G. Megason, “Tissue fluidity mediates a trade-off between the speed and accuracy of multicellular patterning by cell sorting,” Biophysical Journal, vol. 124, no. 23, pp. 4157–4175, 2025.

[81] F. Xiong, A. R. Tentner, P. Huang, A. Gelas, K. R. Mosaliganti, L. Souhait, N. Rannou, I. A. Swinburne, N. D. Obholzer, P. D. Cowgill, et al., “Specified neural progenitors sort to form sharp domains after noisy shh signaling,” Cell, vol. 153, no. 3, pp. 550–561, 2013.

[82] T. Y.-C. Tsai, M. Sikora, P. Xia, T. Colak-Champollion, H. Knaut, C.-P. Heisenberg, and S. G. Megason, “An adhesion code ensures robust pattern formation during tissue morphogenesis,” Science, vol. 370, no. 6512, pp. 113–116, 2020.

[83] C. Collinet and T. Lecuit, “Programmed and self-organized flow of information during morphogenesis,” Nature Reviews Molecular Cell Biology, vol. 22, no. 4, pp. 245–265, 2021.

[84] Y. T. Loo, J. Chen, R. Harrison, T. Rito, S. Theis, G. Charras, J. Briscoe, and T. E. Saunders, “Boundary constraints can determine pattern emergence,” bioRxiv, pp. 2025–07, 2025.

[85] M. S. Steinberg, “Does differential adhesion govern self-assembly processes in histogenesis? equilibrium configurations and the emergence of a hierarchy among populations of embryonic cells,” Journal of Experimental Zoology, vol. 173, no. 4, pp. 395–433, 1970.

[86] F. Graner and J. A. Glazier, “Simulation of biological cell sorting using a two-dimensional extended potts model,” Physical review letters, vol. 69, no. 13, p. 2013, 1992.

[87] T. Y.-C. Tsai, R. M. Garner, and S. G. Megason, “Adhesion-based self-organization in tissue patterning,” Annual review of cell and developmental biology, vol. 38, no. 1, pp. 349–374, 2022.

[88] S. Watanabe, “Information theoretical analysis of multivariate correlation,” IBM Journal of research and development, vol. 4, no. 1, pp. 66–82, 1960.

[89] A. C. Y. Zhang, P. M. Hoyos, D. Brückner, and G. Tkačik, “Nonlocal decoding of positional and correlational information during development,” arXiv preprint arXiv:2512.20536, 2025.

[90] S. A. Newman and W. D. Comper, “‘generic’physical mechanisms of morphogenesis and pattern formation,” Development, vol. 110, no. 1, pp. 1–18, 1990.

[91] A. Hallou, R. He, B. D. Simons, and B. Dumitrascu, “A computational pipeline for spatial mechano-transcriptomics,” Nature methods, pp. 1–14, 2025.

[92] T. G. Andrews, W. Pönisch, E. K. Paluch, B. J. Steventon, and E. Benito-Gutierrez, “Single-cell morphometrics reveals ancestral principles of notochord development,” Development, vol. 148, no. 16, p. dev199430, 2021.

[93] W. Hur, A. Mukherjee, L. Hayden, Z. Lu, A. Chao, N. P. Mitchell, S. J. Streichan, M. Vergassola, and S. Di Talia, “Topological interactions drive the first fate decision in the drosophila embryo,” Nature Physics, pp. 1–12, 2025.

[94] D. Marr, Vision: A computational investigation into the human representation and processing of visual information. MIT press, 2010.

[95] S. R. Quake, “The cellular dogma,” Cell, vol. 187, no. 23, pp. 6421–6423, 2024.

[96] K. J. Mitchell and N. Cheney, “The genomic code: The genome instantiates a generative model of the organism,” Trends in Genetics, vol. 41, no. 6, pp. 462–479, 2025.

[97] S. Wolfram, “Statistical mechanics of cellular automata,” Reviews of modern physics, vol. 55, no. 3, p. 601, 1983.

[98] M. B. Elowitz, A. J. Levine, E. D. Siggia, and P. S. Swain, “Stochastic gene expression in a single cell,” Science, vol. 297, no. 5584, pp. 1183–1186, 2002.

[99] A. Treves and S. Panzeri, “The upward bias in measures of information derived from limited data samples,” Neural Computation, vol. 7, no. 2, pp. 399–407, 1995.

[100] S. Panzeri and A. Treves, “Analytical estimates of limited sampling biases in different information measures,” Network: Computation in neural systems, vol. 7, no. 1, p. 87, 1996.

[101] S. P. Strong, R. Koberle, R. R. D. R. Van Steveninck, and W. Bialek, “Entropy and information in neural spike trains,” Physical review letters, vol. 80, no. 1, p. 197, 1998.

[102] M. Serra and G. Haller, “Objective eulerian coherent structures,” Chaos: An Interdisciplinary Journal of Nonlinear Science, vol. 26, no. 5, 2016.

[103] E. Tang and R. Golestanian, “Quantifying configurational information for a stochastic particle in a flow-field,” New Journal of Physics, vol. 22, no. 8, p. 083060, 2020.

[104] L. Cocconi, Y. Shi, and A. Vilfan, “Information-optimal mixing at low reynolds number,” Physical Review Letters, vol. 135, no. 3, p. 037101, 2025.

[105] A. M. Plum and M. Serra, “Dynamical systems of fate and form in development,” in Seminars in Cell & Developmental Biology, vol. 172, p. 103620, Elsevier, 2025.

[106] G. Gómez-Herrero, W. Wu, K. Rutanen, M. C. Soriano, G. Pipa, and R. Vicente, “Assessing coupling dynamics from an ensemble of time series,” Entropy, vol. 17, no. 4, pp. 1958–1970, 2015.

[107] C. W. Granger, “Investigating causal relations by econometric models and cross-spectral methods,” Econometrica: journal of the Econometric Society, pp. 424–438, 1969.

[108] L. Barnett, A. B. Barrett, and A. K. Seth, “Granger causality and transfer entropy are equivalent for gaussian variables,” Physical review letters, vol. 103, no. 23, p. 238701, 2009.

[109] R. G. James, N. Barnett, and J. P. Crutchfield, “Information flows? a critique of transfer entropies,” Physical review letters, vol. 116, no. 23, p. 238701, 2016.

[110] K.-C. Liang and X. Wang, “Gene regulatory network reconstruction using conditional mutual information,” EURASIP Journal on Bioinformatics and Systems Biology, vol. 2008, no. 1, p. 253894, 2008.

[111] P. L. Williams and R. D. Beer, “Nonnegative decomposition of multivariate information,” arXiv preprint arXiv:1004.2515, 2010.

[112] A. J. Gutknecht, A. Makkeh, and M. Wibral, “From babel to boole: The logical organization of information decompositions,” Proceedings of the Royal Society A: Mathematical, Physical and Engineering Sciences, vol. 481, no. 2310, 2025.

[113] K. Kawasaki, “Diffusion constants near the critical point for time-dependent ising models. i,” Physical Review, vol. 145, no. 1, p. 224, 1966.

